# Divergent effects of pathological α-synuclein truncations and mutations on phase separation

**DOI:** 10.1101/2024.11.18.624073

**Authors:** Soumik Ray, Cecilia Chiodaroli, Azad Farzadfard, Antonin Kunka, Katharina Helga Schott, Sophie Hertel, Kristina Mojtic, Louise Kjær Klausen, Céline Galvagnion, Alexander K. Buell

**Affiliations:** Department of Biotechnology and Biomedicine, Technical University of Denmark, Søltofts Plads, Building 227, 2800, Kgs. Lyngby, Denmark; Department of Drug Design and Pharmacology, Faculty of Health and Medical Sciences, University of Copenhagen, Universitetsparken 2, 2100 Copenhagen, Denmark

## Abstract

Phase separated condensates can accelerate α-synuclein (α-Syn) amyloid fibril formation implicated in Parkinson’s disease pathogenesis. The effects of pathological modifications, i.e., truncations and familial mutations on the thermodynamics, material properties, and the extent of amyloid aggregation within α-Syn condensates remain elusive. Here, we quantitatively demonstrate that terminal truncations significantly alter α-Syn phase separation, while familial mutations impart minimal effects. Spontaneous sol-gel phase transitions of the truncated α-Syn variants could give rise to amyloid fibrils almost instantly within condensates, suggesting similarities between molecular interactions driving both processes. Extending our study to model coacervate and condensate systems where α-Syn acts as a client, we find α-Syn can dissolve coacervates and form Pickering clusters on condensate surfaces—regulating their size. Additionally, the C-terminal region of α-Syn modulates nucleic acid sequestration within condensates. Together, our findings reveal diverse effects of α-Syn modifications on phase separation, both in pathological and physiological contexts.

**GRAPHICAL ABSTRACT:** 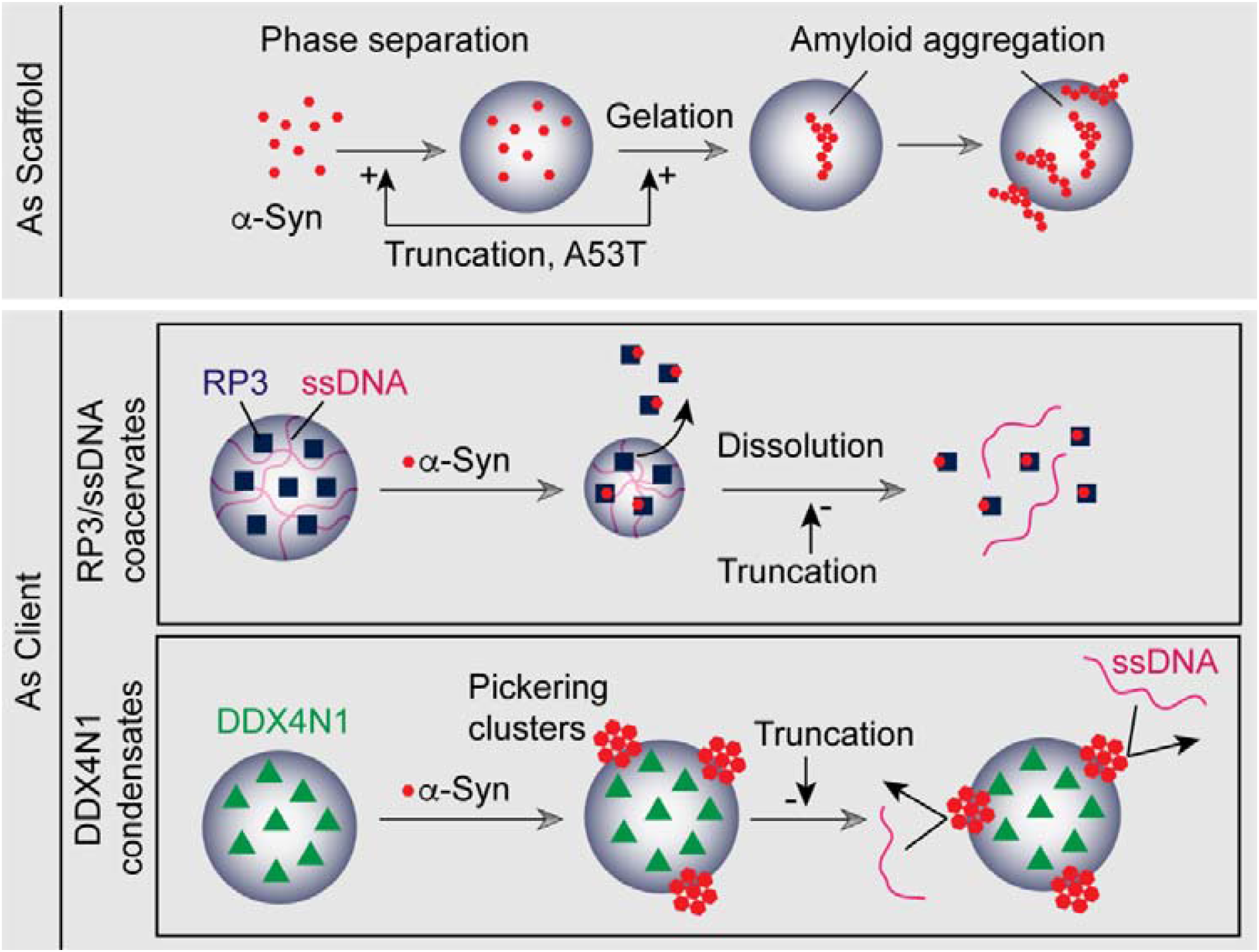

## INTRODUCTION

Intrinsically disordered proteins (IDPs) are prone to interact and assemble into higher-order complexes due to their multivalent nature^1^. One of the most important ways they can assemble is by forming dynamic, reversible membraneless condensates via phase separation^2,3^—regulating many processes involved in the Central Dogma of a living cell^4-7^. Many biomolecules (e.g., proteins and nucleic acids) can sequester within a given condensate in cells—transiently achieving very high local concentrations of individual species that are thought to be able to enhance and compartmentalize biochemical processes^7-13^. However, disease related mutations and post-translational modifications of IDPs often induce rapid formation of amyloid fibrils within condensates, leading to deleterious outcomes^5,14-18^. Apart from potential cytotoxicity of fibrils themselves, aggregated condensates may also irreversibly trap essential cellular components—disrupting a myriad of physiological processes. A growing body of evidence suggests that these processes are linked to the pathogenesis of neurodegenerative diseases such as Amyotrophic Lateral Sclerosis (ALS), Frontotemporal Dementia (FTD), Alzheimer’s (AD), and Parkinson’s disease (PD)—mediated by IDPs like FUS, hnRNPA1, TDP-43, tau, and α-synuclein (α-Syn)^19-26^. To understand the molecular intricacies of neurodegeneration under the lens of phase separation, it is of utmost importance to study how IDP condensates are affected by disease relevant modifications. FUS, hnRNPA1, TDP-43, and tau have been characterized in this regard^19,27-34^. However, the precise effects of pathological α-Syn modifications (associated with PD) remain unclear. Phase separation of α-Syn is particularly intriguing because condensate maturation (via ripening/fusion) concomitant with amyloid aggregation occurs over very slow timescales (hours to days, depending on solution conditions^26^). This slow phase transition can be advantageous in temporally resolving the underlying molecular processes. Despite this potential and relevance, no study have investigated the effects of pathological α-Syn modifications on phase separation quantitatively, with the majority of investigations remaining primarily qualitative in nature^35-37^.

In this body of work, we employ in-house developed, quantitative phase separation assays^38,39^ alongside state-of-the-art biophysical techniques, to characterize α-Syn phase separation with a focus on its thermodynamics, material properties, and amyloid aggregation. We compare these effects among terminal truncations and familial point mutations, and show that truncations impart the most significant changes to both the thermodynamics and material properties of α-Syn condensates. While full-length variants undergo canonical liquid-liquid phase separation, the truncated variants exhibit instantaneous sol-gel phase transitions. Intriguingly, regardless of their gelation timescales (from seconds to hours), all variants form elongation/templating competent amyloid fibrils within the dense phase. In parallel, we also discover unique functions of α-Syn as a ‘client’ molecule within model coacervate (RP3/ssDNA^38^) and condensate (DDX4N1^38,40^) systems. The RP3/ssDNA coacervates and DDX4N1 condensates, in our case, represent two major models of phase separation in cells— driven by electrostatic interactions and hydrophobic/cation-π interactions, respectively^9^. We show that by either partitioning within coacervates, or adsorbing on condensate surfaces as sub-micron scale Pickering clusters, α-Syn can regulate nucleic acid sequestration within these compartments. Our experiments reveal that the negatively charged C-terminal of α-Syn is a primary modulator of these effects, as these effects become less pronounced in the C-terminally truncated variants.

## RESULTS

### Pathological modifications alter the thermodynamics of α-Syn phase separation

We employed Taylor dispersion-induced phase separation (TDIPS^39^), a fully automated capillary-based technique we developed on the FIDA1 instrument, to chart phase diagrams of α-Syn variants as a function of NaCl and protein concentration. Here, we quantified the extent of phase separation by multiplying the number of detected spikes in fluorescence intensity corresponding to condensates (n) with the average intensity of the spikes (I_avg_) (Fig. 1a, left, Supplementary Fig. 1). The wild-type (WT) and familial mutants (A30P, H50Q, G51D and A53T) did not phase separate in the absence of NaCl— suggesting Debye screening of unfavourable electrostatic interactions by NaCl is required for demixing to occur. Interestingly, apart from a broader phase diagram, i.e., formation of condensates at lower protein and NaCl concentrations, C-terminally truncated (residues 1-125, ‘ΔC1-125’) and both N/C-terminally truncated (residues 30-110, ‘Core’) α-Syn showed an opposite behavior: these variants phase separated more strongly in the absence of NaCl^35^ (Fig. 1a-left). This was further verified using confocal microscopy (Fig. 1a, right). The dilute phase concentration (C_dil_) is a crucial parameter to determine the thermodynamic stability of condensates. This is because *C*_*dil*_ D< *e*^−χ^, and the Flory parameter (χ) is a measure of the interaction strength between protein molecules undergoing phase separation, relative to solvent interactions^41^. We measured C_dil_ of α-Syn variants as a function of NaCl concentration using UV absorption and Capillary flow experiments (Capflex^38^)— the latter being another method that we developed on the FIDA1 specifically to quantify the dilute phase concentrations (Supplementary Fig. 1). We found a systematic decrease in C_dil_ with increasing NaCl concentration for WT and familial α-Syn mutants (A30P, H50Q, G51D and A53T), albeit to varying extents (Fig. 1b, left, Supplementary Fig. 2). At 250 mM NaCl, significant differences in C_dil_ were observed for H50Q and A53T, with average values of 76 μM and 80 μM respectively, compared to 125 μM for WT (Fig. 1b, right). In line with TDIPS, C_dil_ of truncated α-Syn variants exhibited an inverted relation with NaCl concentration—phase separating more strongly with decreasing ionic strength. At 0 mM NaCl, C_dil_ scaled well with the degree of truncation, being the lowest for Core (46 μM), followed by ΔC1-110 (80 μM) and ΔC1-125 (135 μM) (Fig. 1c, Supplementary Fig. 2). From these observations, we hypothesized that the negatively charged (acidic) amino acids (abundant in the C-terminal region) were responsible for the inversion of salt dependence. To demonstrate this further, we measured C_dil_ of a recombinant α-Syn variant (C_rev_), where five C-terminal acidic residues were substituted with lysine^26^. Akin to truncated variants, phase separation of C_rev_ was also electrostatically favourable with a lowest C_dil_ of 145 μM at 0 mM NaCl (Fig. 1c). Subsequently, we measured C_dil_ of WT and C-terminal truncated/modified variants across a pH range of 5.6-7.8. Our data showed that the formal net charge of α-Syn variants (theoretically derived) correlated well with the extent of phase separation at different pH values (Fig. 1d-e)—establishing the importance of the charged C-terminal tail in modulating α-Syn phase separation.

**Figure 1:**
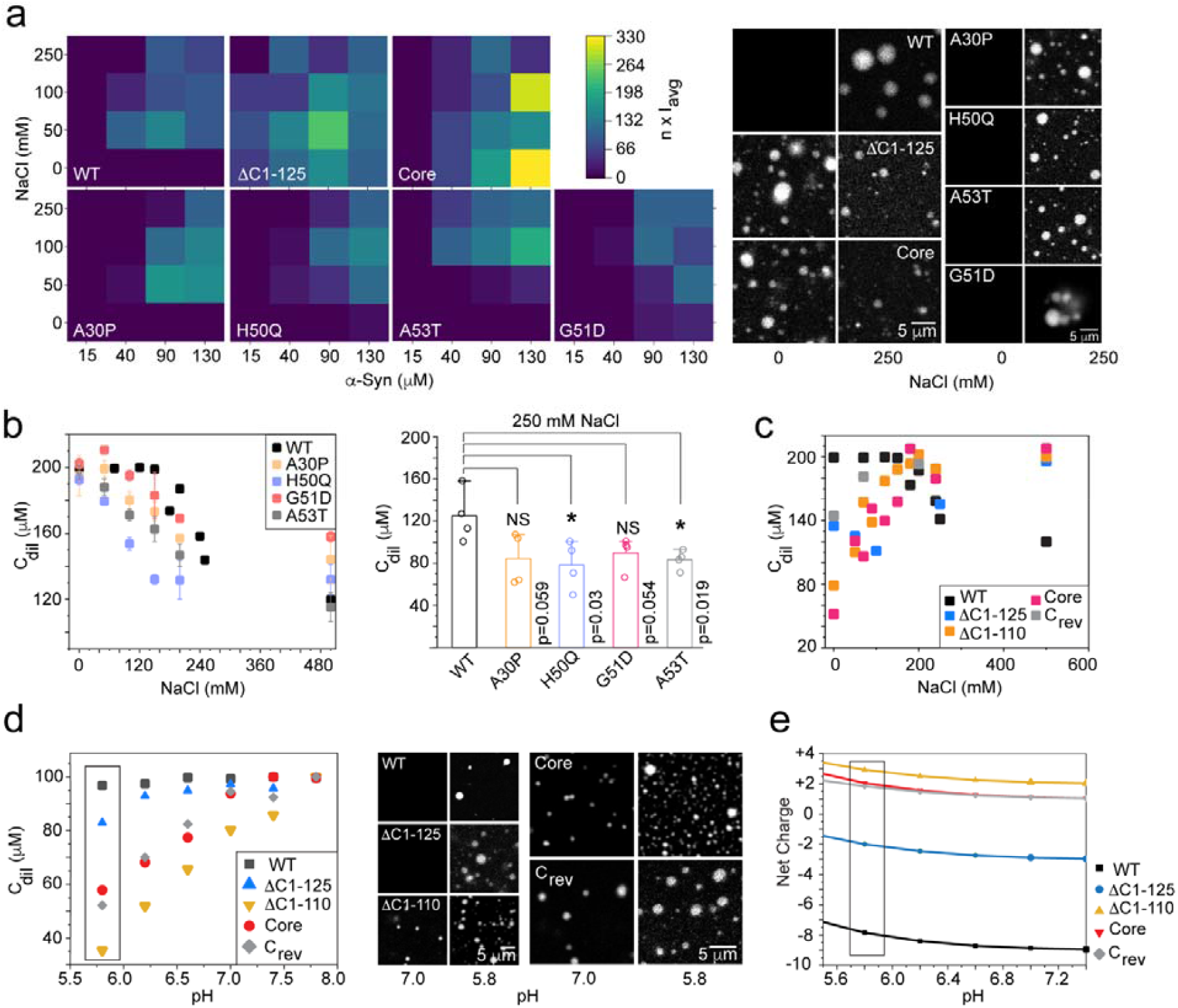
Thermodynamics of phase separation of α-Syn variants: **a**. (Left) Full phase diagram of WT and variant (ΔC1-125, Core, A30P, H50Q, A53T, G51D) α-Syn as a function of NaCl concentration. Here, n=number of condensates detected, and I_avg_=average intensity of fluorescence spikes/condensates. (Right) Representative confocal microscopy images of the said proteins under 0 mM and 250 mM NaCl. The experiments are performed with 25% (w/v) PEG-8000, in 20 mM sodium phosphate buffer, and at 25°C. **b**. (Left) C_dil_ as a function of NaCl concentration for WT and familial (A30P, H50Q, G51D, A53T) α-Syn mutants. (Right) Comparative C_dil_ of WT and familial α-Syn mutants at a fixed (250 mM) NaCl concentration. Values represent mean±SD for n=3 independent experiments. Statistical significance is calculated with one-way ANOVA at a 95% confidence interval. The p-values are shown in the figure. **c**. C_dil_ as a function of NaCl concentration for WT and terminally truncated/altered α-Syn variants (ΔC1-125, ΔC1-110, Core, C_rev_). The experiments (**b-c**) are performed at C_t_=200 μM with 20% (w/v) PEG-8000, in 20 mM sodium phosphate buffer, pH 7.4, and at 25°C. **d**. (Left) C_dil_ as a function of pH for WT and terminally truncated/altered α-Syn variants (ΔC1-125, ΔC1-110, Core, C_rev_). The experiments are performed at C_t_=100 μM with 15% (w/v) PEG-8000, in 20 mM sodium phosphate buffer, and at 25°C. (Right) Representative confocal microscopy images of the said proteins at pH 5.8 and 7.4. **e**. Theoretically derived net charge of WT and terminally truncated/altered α-Syn variants (ΔC1-125, ΔC1-110, Core, and C_rev_). The rectangles in **d-e** highlight similar trends between experimentally derived C_dil_ and the net charge of α-Syn variants.

**Figure 2:**
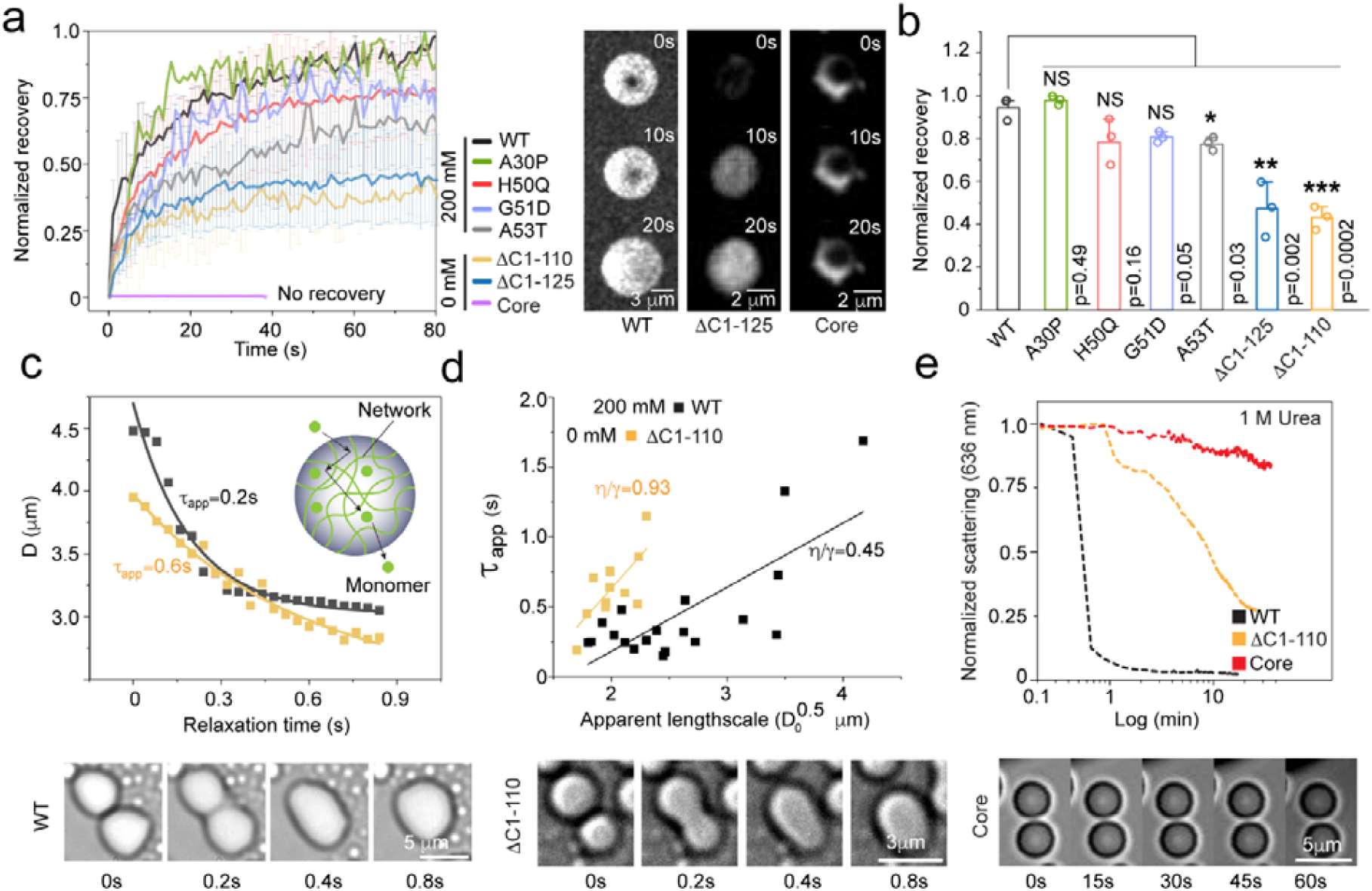
Condensate material properties of α-Syn variants: **a**. (Left) Normalized FRAP recovery of WT and variant α-Syn condensates under optimal conditions (immediately after formation). Meaning that for WT and familial mutants, the condensates are formed at 250 mM NaCl, and truncated variant condensates are formed at 0 mM NaCl. (Right) Representative confocal microscopy images of WT, ΔC1-125 and Core condensates at 0s, 10s and 20s post bleach. **b**. Statistical analysis of the normalized recovery of variant α-Syn condensates compared to the WT. Values represent mean±SD for n=3 independent experiments. Significance is calculated with one-way ANOVA at a 95% confidence interval. The p-values are shown in the figure. C_t_=200 μM for each protein. The experiments are performed with 20% (w/v) PEG-8000, in 20 mM sodium phosphate buffer, pH 7.4, and at 25°C. **c**. Distance (D) as a function of relaxation time is plotted for two representative fusion events detected for WT and ΔC1-110. **d**. τ_app_ values as a function of D_0_ are plotted for WT and ΔC1-110. The value of the slopes (corresponding to υ_app_=η/γ) obtained from linear regression are indicated in the figure. (**c-d**) (Bottom) Representative bright-field (BF) microscopy images showing fusion events upon contact for WT and ΔC1-110 condensates. Note that despite smaller sizes (scale bars), ΔC1-110 condensates fuse and relax with similar timescales as larger WT condensates. Core condensates (right) do not fuse upon contact even after 1 min. **e**. 636 nm light scattering showing WT condensates dissolve upon addition of 1 M urea within 1 min, while ΔC1-110 and Core condensates persist. The persistence times corroborates with the FRAP/fusion data.

It is important to note here that in our Capflex experiments, we used 50 nM Alexa488-140C-α-Syn as a fluorescent probe to quantify the C_dil_ for all α-Syn variants. Remarkably, the concentration of the labeled α-Syn in the dilute phase accurately reflected the C_dil_ of all purely unlabeled variant counterparts (Supplementary Fig. 2), regardless of their differences in phase separation behavior. It was particularly intriguing that the labeled α-Syn could also be used as a probe for the truncated variants, which exhibit a reversed electrostatic behaviour. This could be due to their large difference in concentration, with the unlabeled α-Syn being 10^4^ times more abundant (total concentration, C_t_=200 μM). The labeled α-Syn, possessing all relevant moieties for interactions (since it is full-length), likely experiences dominant interactions based on the characteristics of the unlabeled counterpart, thus dominating other interactions that might otherwise lead to different phase separation behaviour. To test this further, we verified the partitioning of Alexa488-140C-α-Syn in a co-phase separating system with both WT and C_rev_ α-Syn. Under our experimental conditions (pH 6.0, C_t_=25 μM), C_rev_ phase separates more than WT with a significantly lower C_dil_ (Supplementary Fig. 2). The idea was to systematically increase the fraction of the WT monomers relative to the C_rev_ (keeping the total α-Syn monomer concentration constant) to observe whether the labeled α-Syn could follow the change in the C_dil_ as the WT fraction increased. Indeed, the baseline fluorescence (representing C_dil_) increased systematically as a function of the WT fraction—indicating that the labeled α-Syn represented the overall phase behavior of the mixed system (Supplementary Fig. 2). We also noted that the increase in the baseline fluorescence did not follow a linear relationship with the WT fraction, suggesting that the interactions were not linearly dependent on the proportion of WT in the mixture (Supplementary Fig. 2). This reflects the fact that every molecule inside a condensate interacts with more than one other molecule.

### Condensates formed by truncated α-Syn variants show more gel-like material properties

We employed fluorescence recovery after photo bleaching (FRAP^42^) to examine the translational dynamics of molecules within WT and variant α-Syn condensates. Note that all data represent spontaneous demixing without any time delay and under conditions where maximum phase separation is observed for each variant (see methods). Condensates formed by WT and familial mutants recovered well (∼90%), except for A53T (∼75%) (Fig. 2a-b). Interestingly, ΔC1-110 and ΔC1-125 exhibited significantly decreased fluorescence recovery (∼50%), while fluorescence from Core condensates did not recover at all—suggesting a sol-gel phase transition with very fast kinetics (Fig. 2a-b). Two-dimensional FRAP alone is not sufficient to infer macroscopic material properties since monomers can diffuse freely even within a network architecture inside condensates (Fig. 2c, inset)^43^. To verify whether the observed translational dynamics were in agreement with the macro-viscosity (*η*) of condensates, we measured the relaxation kinetics of condensates during fusion events^44,45^ (Fig. 2c-d, Supplementary Fig. 3). We restricted subsequent experiments to only the WT, ΔC1-110, and Core which showed significant differences in FRAP. Tens of fusing condensate pairs that had similar sizes were chosen for both WT and ΔC1-110. In a representative case, monoexponential fitting of the distance (D) as a function of relaxation time (t) provided us with a time constant: τ_app_=0.2s for WT with a D_0_=4.50 μm (distance at the beginning of fusion). Despite a lower D_0_=4.0 μm (smaller condensates), τ_app_=0.6s was considerably higher for ΔC1-110 (Fig. 2c). Plotting τ_app_ as a function of √D_0_ (related to the lengthscale of fusion, see method) yielded the apparent inverse capillary velocity: 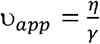, which was ∼2 times greater for ΔC1-110 (υ_app_∼0.93 sμm^-1^) compared to WT (υ_app_∼0.45 sμm^-1^) (Fig. 2c, right, Supplementary Fig. 3). To note, υ_app_ does not directly translate to *η*, since the surface tension (γ) of condensates can be different between the two proteins. The scatter of the WT data arose from some of the condensate pairs in contact with the glass slide—experiencing an additional force imparted by the surface that slowed down fusion, and prevented complete relaxation. Therefore, the determined value of υ_app_ is likely to be an overestimate for WT. Our simplified analyses showed that the overall material properties were significantly different—ΔC1-110 condensates being more viscous compared to those formed by the WT, while Core condensates were essentially solid-like and showed no fusion upon contact (Fig. 2d). These results were further supported by light scattering-based condensate dissolution experiments in presence of 1 M urea. Here, ΔC1-110 and Core condensates showed remarkable resistance against dissolution—remaining stable for tens of minutes, while WT dissolved within 1 minute (Fig. 2e).

**Figure 3:**
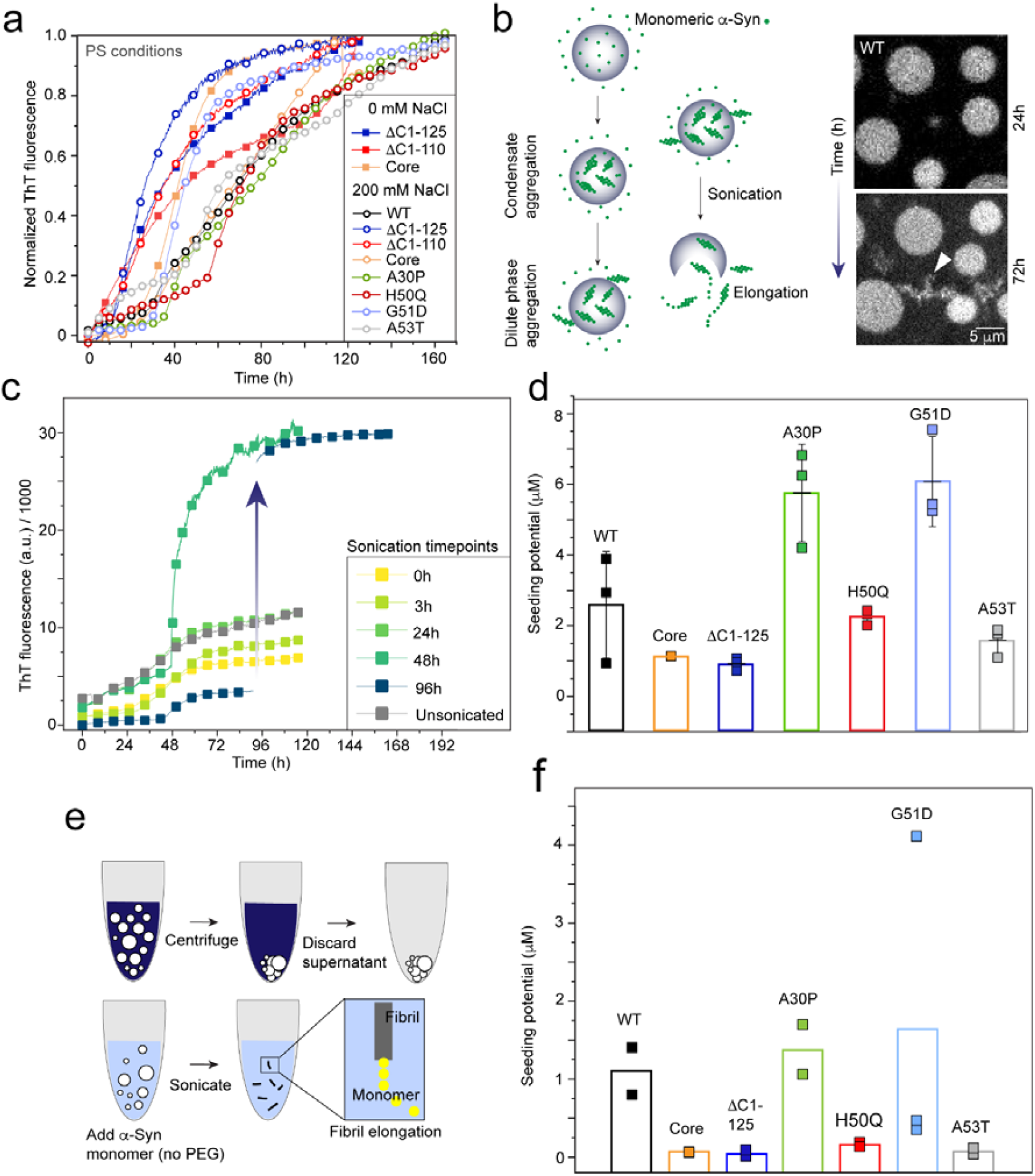
Condensate-mediated amyloid aggregation of α-Syn variants: **a**. Ensemble ThT aggregation kinetics of α-Syn variants at 0 mM and 250 mM NaCl. All samples that are phase separated show aggregation. The experiments are performed under quiescent conditions, with 20% (w/v) PEG-8000, in 20 mM sodium phosphate buffer, pH 7.4, and at 25°C. **b**. (Left) a schematic depicting the aggregation landscape in α-Syn condensate solutions undergoing dense phase aggregation followed by dilute phase aggregation during ageing. Sonicating α-Syn condensates post dense phase aggregation (and before majority of dilute phase aggregation) results in release of amyloid fibrils (if any) from the condensates. The exposed fibrils can then elongate in the presence of dilute phase monomers. (Right) Representative confocal fluorescence microscopy images of WT α-Syn condensates at 24h and 72h. At 72h, the dilute phase monomers start to aggregate/localize on condensate surfaces (marked with a white triangular pointer). 100 nM Alexa488-140C-α-Syn is used as a fluorescent reporter. **c**. ThT fluorescence of 200 μM WT α-Syn condensate solution sonicated at different timepoints. Note that fibril elongation is only observed when the condensates are sonicated after 48h of incubation. The blue arrow indicates a sudden increase in ThT fluorescence when the sample is sonicated at 96h. The experiments are performed under quiescent conditions, with 20% (w/v) PEG-8000, in 20 mM sodium phosphate buffer, pH 7.4, and at 25°C. **d**. Seeding potential for WT and variant α-Syn condensates sonicated at 48h (full-length) and 0h (truncated), under optimal phase separating conditions. Values represent mean±SD for n=3 independent experiments. **e**. Schematic showing the experimental strategy to calculate seeding potential of condensates by isolating them by centrifugation. **f**. Seeding potential for WT and variant α-Syn condensates isolated by centrifugation and subsequently sonicated at 48h (full-length) and 0h (truncated), under optimal phase separating conditions. Datapoints and average are shown from n=2 (n=3 for G51D) independent experiments.

### Pathological modifications lead to different extents of amyloid fibril formation via phase separation

We next set out to quantify amyloid aggregation within variant α-Syn condensates. We performed bulk Thioflavin-T (ThT) fluorescence-based kinetics experiments under quiescent conditions and at 25°C (Fig. 3a). Under these conditions, *de novo* amyloid aggregation independent of phase separation could be minimized (Supplementary Fig. 4). ThT fluorescence of condensate solutions of WT and familial mutants showed a slow increase until ∼50h, followed by a rapid rise and saturation (Fig. 3a). Aggregation in condensate solutions of ΔC1-125, ΔC1-110 and Core were considerably faster, reaching saturation within ∼50h (Fig. 3a)—a finding in agreement with the FRAP and fusion data. However, we argued that such bulk measurements might not necessarily reflect the extent of amyloid aggregation within condensates, as solid-like condensates without containing any amyloid fibrils could still provide high-affinity surfaces for the dilute phase monomers to initiate amyloid aggregation (akin to lipid vesicle/nanoparticle induced amyloid aggregation^46,47^). Transmission electron microscopy (TEM) imaging showed presence of fibrillar aggregates after 7 days of incubation for all phase separated variants (Supplementary Fig. 4), suggesting that regardless of the pathway, condensates ultimately gave rise to amyloid fibrils. To determine the relative amounts of amyloid fibrils formed within condensates during ageing, we sonicated phase separated solutions at specific time intervals. We aimed to identify precise timepoints when amyloid aggregates have formed within condensates, but the majority of the dilute phase monomers has not yet aggregated. Fragmenting such condensates using sonication will expose amyloid fibrils to the bulk solution containing monomeric α-Syn, and immediately initiate an enhanced fibril elongation reaction—rates of which can be used to approximately estimate the original fibril concentration in monomer equivalent (Fig. 3b-c). We ensured that the amplitude and frequency of sonication could disrupt the aggregated condensates, and that sonication itself does not induce α-Syn aggregation (Supplementary Fig. 4). Condensate fragmentation can create more surface area, but the resulting aggregation kinetics (after sonication) would be dominated by the elongation mechanism in presence of fibrils. For WT, sonication up to 24h showed no change in ThT fluorescence compared to an unsonicated control (Fig. 3c). Strikingly, sonication at 48h resulted in a rapid increase in ThT fluorescence indicative of elongation, followed by saturation (Fig. 3c). Sonication at 96h led to an immediate increase (∼factor of 3) of ThT intensity, however with no signs of fibril elongation (Fig. 3c, arrow). Our data suggested that elongation competent amyloid aggregates within WT condensates appear at ∼48h. Between 48 and 96 hours, amyloid aggregation progresses within the dense phase and extends to the monomers in the dilute phase, which are in dynamic exchange with the condensates (Fig. 3b, right). Beyond 96h, the limited availability of monomers precludes further elongation. The increased ThT intensity after sonication at this late time point presumably arises due to the exposure of the amyloid fibrils to ThT after condensate disruption. We chose the 48h mark to compare elongation profiles of familial mutant α-Syn condensates using the same strategy. Since the truncated α-Syn condensates were gel-like from the beginning (therefore, could potentially catalyze surface nucleated amyloid aggregation at 0h), we sonicated them at 0h to ensure the majority of the fibrils, if any were to be detected, had originated within condensates (Supplementary Fig. 5). The elongation rates were calculated using linear regression of ThT fluorescence values obtained for several (4-5) hours post-sonication (Supplementary Fig. 5). We acknowledge that the available monomer concentration [m] was different for each phase separating system having different C_dil_. However, the elongation rate constant (k_+_) becomes independent of [m] above a threshold concentration of several tens of micromolar, in most cases^48,49^. For 2.5 μM α-Syn fibril seeds at pH 7.4. We calculated it to be ∼30 μM (Supplementary Fig. 5), which was less than the measured C_dil_ value (Fig. 1). Therefore, we assumed that differences in [m] imparted no significant changes to the measured k_+_. Using calibration curves derived from the 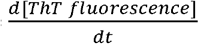 values of known _dt_ concentrations of pre-made, sonicated fibril seeds at fixed (30 μM) monomer concentration (Supplementary Fig. 5), we estimated fibril concentrations within aggregated condensates for each α-Syn variant (Fig. 3d). According to our calculations, out of a C_t_ of 200 μM, aggregated WT condensates contained ∼3 μM elongation competent amyloid fibril seeds. To note, this assumes that the fibrils within condensates are fragmented by sonication into seeds of lengths comparable to standard seeds used in calibration experiments (∼100 nm). Among familial mutants, A30P and G51D had the highest fibril seed concentrations (∼6 μM), followed by H50Q (∼2 μM), A53T (∼1.5 μM). Remarkably, the truncated mutants also showed presence of elongation competent fibril seeds (∼1 μM) even at 0h. This was surprising as we expected instantaneous gelation of these variants might not predominantly result in amyloid fibril formation within the dense phase, but rather to non-fibrillar structures such as amorphous gels. These observations suggested that the molecular interactions driving phase separation of truncated α-Syn variants might not be very different from the interactions responsible for amyloid fibril formation. We term the estimated fibril seed concentrations ‘*seeding potential* (Fig. 3d)’ since the dominant fibril polymorph formed within the condensates may not exhibit identical elongation kinetics compared to fibrils that are used as seeds for the calibration curves (prepared under shaking, in absence of PEG)^50^. To support our observations, we separated aggregated α-Syn condensates (48h for full-length and 0h for truncated variants) using centrifugation and measured the seeding potential of the isolated condensates (Fig. 3e-f). The advantage of this approach was that it did not require the calibration experiments to be performed in the presence of PEG, which can induce undesired phase separation (see methods). Here, the relative differences in seeding potential among the variants remained very similar to our previous experiment (Fig. 3f). However, we noted that the seeding potentials were substantially decreased across all α-Syn variants, and also observed considerable differences between independent experiments for G51D—both of which could be due to the limitations in centrifuging the dense phase entirely in viscous PEG solutions. We elaborate on these technical aspects further in the method section.

**Figure 4.**
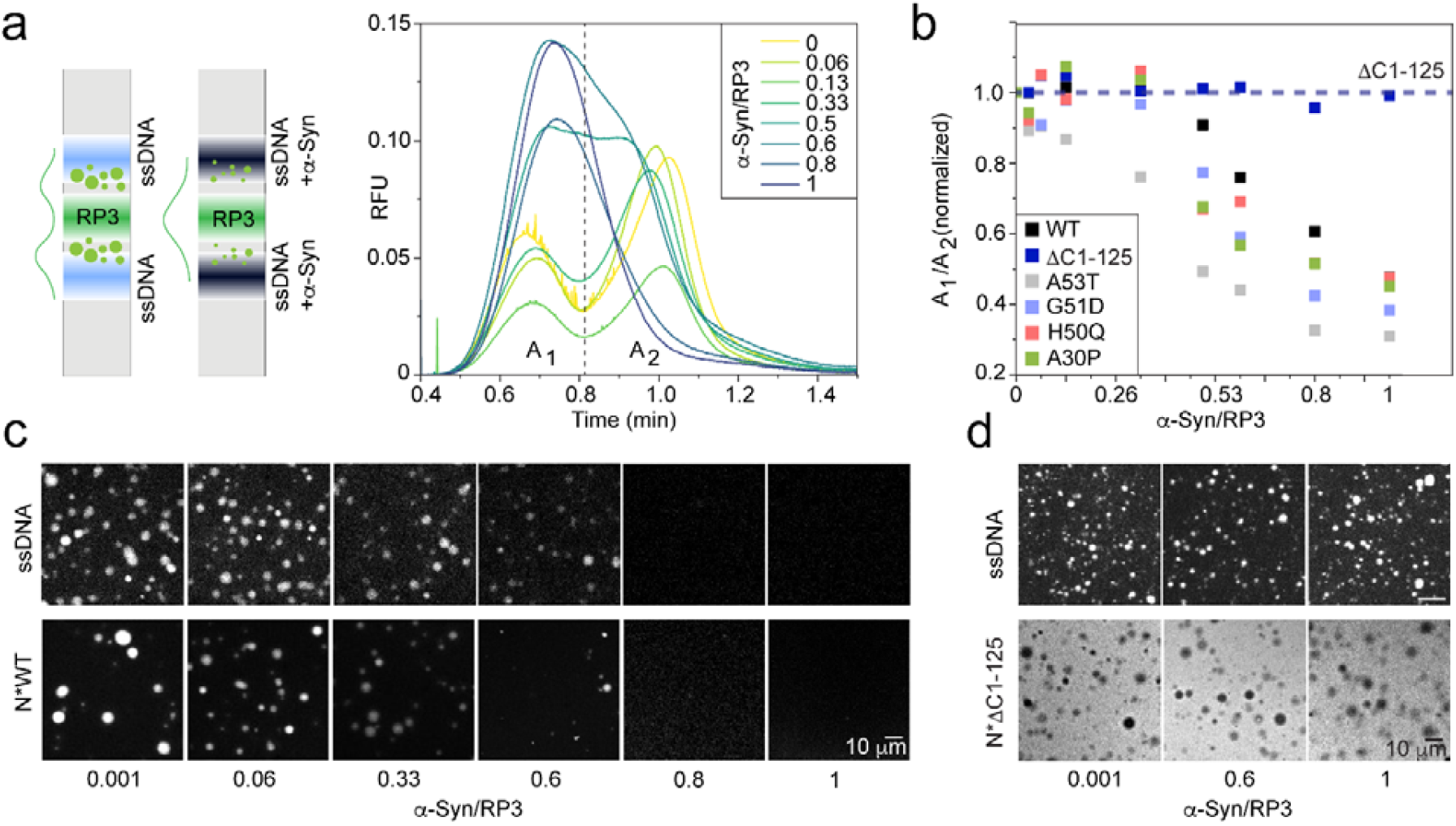
α-Syn as client for electrostatic coacervates: **a.** (Left) A triple-plug TDIPS strategy to investigate the effects of α-Syn on RP3/ssDNA coacervate system. (Right) In the absence of α-Syn, RP3 and ssDNA form coacervates, which is detected as splitting (mass transfer) of an otherwise Gaussian profile. Increasing α-Syn concentrations in the ssDNA plugs dissolve coacervates in a dose-dependent manner. **b**. The ratio of the area under the two peaks (A_1_/A_2_) when coacervates form are plotted as a function of WT and variant α-Syn concentrations (shown as a stoichiometric ratio relative to RP3). The dashed line highlights that ΔC1-125 cannot dissolve coacervates. **c**. Representative confocal microscopy images of RP3/ssDNA coacervates as a function of WT α-Syn. The top and bottom panels show experiments where the fluorophore is Alexa488-ssDNA and N-terminally labeled Alexa488-WT-α-Syn, respectively. **d**. Representative confocal microscopy images of RP3/ssDNA coacervates as a function of ΔC1-125 α-Syn. The top and bottom panels show experiments where the fluorophore is Alexa488-ssDNA and N-terminally labeled Alexa488-ΔC1-125-α-Syn, respectively. For α-Syn/RP3=0.001, only labeled (without unlabeled) α-Syn is added to the reaction mixture. The experiments (**c-d**) are performed with 200 μM RP3, 20 μM ssDNA, in 10 mM Tris-HCl, pH 8.0, 150 mM NaCl, and at 20°C.

**Figure 5.**
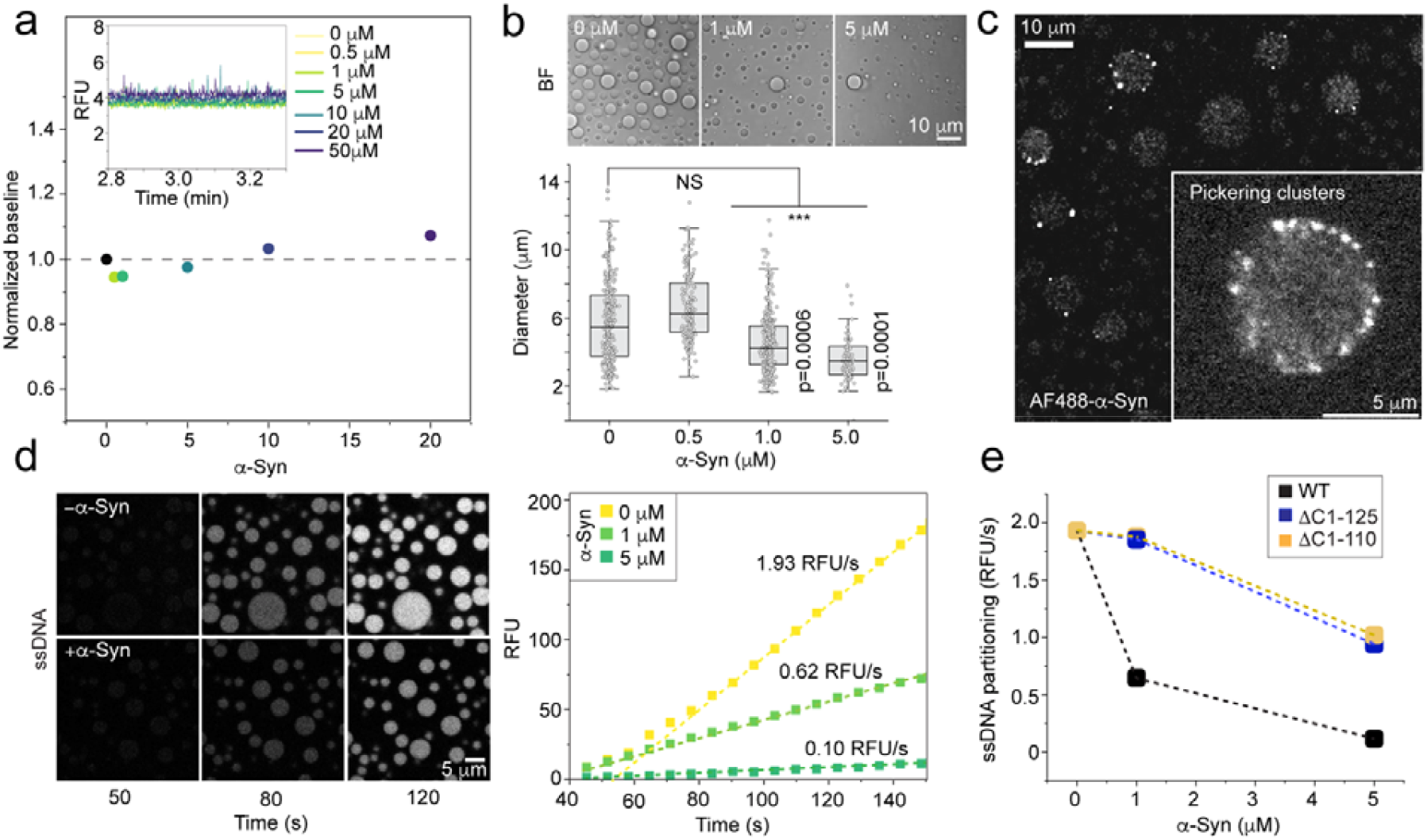
α-Syn as client for DDX4N1 condensates: **a.** Capflex traces (inset) and normalized baseline fluorescence showing increasing α-Syn concentrations have minimal effects on the C_dil_ of DDX4N1 phase separation. **b**. (Top) Representative bright field (BF) microscopy images showing DDX4N1 condensate size decrease as a function of α-Syn concentration. (Bottom) Distribution of DDX4N1 condensate diameter as a function of α-Syn concentration. Statistical significance is calculated with one-way ANOVA at a 95% confidence interval. The p-values are shown in the figure. **c**. Representative confocal microscopy images with 500 nM Alexa-488 labeled 140C-α-Syn showing Pickering clusters of α-Syn on the surface of DDX4N1 condensates. **d**. (Left) Representative time-lapse images showing partitioning of 50 nM Alexa488-ssDNA within DDX4N1 condensates in the absence and presence of 1 μM α-Syn. (Right) DDX4N1 condensate fluorescence intensity (background corrected) during ssDNA partitioning is plotted and fitted with a linear equation. The slopes (relative fluorescence unit (RFU)/s) are indicated in the presence of increasing α-Syn concentrations. The datapoints represent mean values obtained from N=50 condensates. **e**. ssDNA partitioning rates for WT, ΔC1-125, and ΔC1-110 with increasing α-Syn concentrations. The experiments **(a-e)** are performed with 70 μM DDX4N1, in 10 mM sodium phosphate buffer, pH 6.5, 50 mM NaCl, and at 25°C.

Next, to explore the impact of amyloid fibril formation via phase separation on cellular health, we measured the toxicity of these fibrils in human neuronal cells. We performed an MTT assay on undifferentiated SH-SY5Y cells, with 5 μM monomer equivalent concentrations of α-Syn (WT and mutant) condensate solutions that are aged for 7 days (i.e., the majority of the protein is aggregated with very little dilute phase monomer left). Our results revealed that condensate derived fibrils substantially reduced cell viability (70% after 24h, and 60% after 96h) (Supplementary Fig. 6). Surprisingly, among the α-Syn variants analyzed, fibrils derived from the Core α-Syn variant showed no detectable toxicity. This unexpected outcome could be explained by two possible scenarios: 1) Fibrils formed in the presence of core condensates were inherently non-toxic due to their different modes of interaction with cells stemming from the complete lack of the disordered flank regions. 2) Another explanation lies in the unique phase behavior of the Core α-Syn condensates. Core α-Syn condensates accumulate majority of the monomers in the dense-phase, leaving very little dilute phase available (Fig. 1). Despite this, Core α-Syn does not appear to form the same amounts of fibrils within condensates as WT after gelation (Fig. 3), and perhaps the major fraction of the solidified condensates consist of non-fibrillar material. Therefore, the total amount of fibrils that the system can form are potentially lower, resulting in a lower toxicity. This observation underscores how the relative distribution between dense and dilute phases as well as the nature of the dense phase can drastically influence the potential pathogenicity of α-Syn fibrils derived from condensates.

### α-Syn can dissolve electrostatic coacervates

Like many other IDPs, α-Syn can also be sequestered within other physiological coacervates/condensates and regulate their composition and/or stability^51-53^. To explore this aspect for our collection of α-Syn variants, we first studied α-Syn as a ‘client’ molecule in a model coacervate system consisting of a short cationic peptide (RP3: (RRASL)_3_) and a single stranded nucleic acid sequence (ssDNA, 30 nucleotides)^38^. In a TDIPS experiment, we placed an RP3 plug between two ssDNA plugs (with and without α-Syn, see methods) (Fig. 4a, left) to induce RP3/ssDNA coacervation within a capillary. In the absence of α-Syn, the expected Gaussian peak of ssDNA split into two peaks with detectable spikes—indicating mass transfer from the main peak due to phase separation^39^ (Fig. 4a, right). Interestingly, as the α-Syn concentration within the ssDNA plugs increased, these two distinct peaks (and spikes) gradually disappeared, and a single peak (no phase separation) emerged at equimolar α-Syn to RP3 ratios (Fig. 4a, right). This suggested that α-Syn could alter the electrostatic balance and dissolve RP3/ssDNA coacervates. We quantified the coacervate dissolution potential of WT and pathological α-Syn variants by plotting the ratio between the areas under the two peaks, as a function of α-Syn concentration (Fig. 4b, Supplementary Fig. 7). We found that all full-length variants dissolved RP3/ssDNA coacervates at near equimolar ratios (0.8-1) of α-Syn to RP3 (Supplementary Fig. 7). Intriguingly, ΔC1-125 could not dissolve the RP3/ssDNA coacervates—stressing the importance of the negatively charged C-terminal region for this effect (Fig. 4b, Supplementary Fig. 7). In addition, we noted that A53T dissolved coacervates more effectively than the rest of the full-length variants (Fig. 4b)—suggesting additional interactions at the N-terminal region could also be important for the interplay between α-Syn and RP3/ssDNA coacervates. To establish that the C-terminal region acted as a proxy for ssDNA by binding to RP3, we performed parallel confocal microscopy experiments with Alexa488-ssDNA and N-terminally labeled Alexa488-α-Syn (both WT and ΔC1-125). WT partitioned homogeneously within RP3/ssDNA coacervates and dissolved them in a dose-dependent manner, while ΔC1-125 could not (Fig. 4c-d). Moreover, we found that ΔC1-125 remained excluded from the coacervates, which stands in agreement with previous observations^54^. Taken together, the microscopy data fully agreed with the TDIPS experiments, and clearly showed that electrostatic regulation of coacervates was dictated by the interactions between RP3 and the negatively charged C-terminal tail of α-Syn.

### α-Syn acts as Pickering agent for DDX4N1 condensates

Next, we investigated the effects of α-Syn on DDX4N1^38,40^ condensates. Capflex revealed that α-Syn did not have any substantial effects on the thermodynamics of DDX4N1 phase separation (Fig. 5a), as the C_dil_ of DDX4N1 remained unchanged (C_dil_=53 μM, with a C_t_=70 μM) as a function of α-Syn concentration. However, DDX4N1 condensates appeared significantly smaller in the presence of sub-stoichiometric α-Syn concentrations (as low as 1 μM α-Syn, relative to 70 μM DDX4N1) (Fig. 5b). Strikingly, confocal microscopy revealed that α-Syn formed distinct submicron-scale clusters on the surface of DDX4N1 condensates (Fig. 5c). These observations suggested that α-Syn might act as a Pickering agent^55,56^ on DDX4N1 condensates—providing a steric effect that can restrict fusion mediated growth of the condensates. Indeed, DDX4N1 condensate fusion events in the presence of 5 μM α-Syn was substantially reduced in frequency (100 events over 1 min) compared to a no α-Syn control (40 events over 1 min) (Supplementary Fig. 8). Except for Core α-Syn, all other α-Syn variants (including the truncated ΔC1-125) resulted in size reduction of the DDX4N1 condensates (Supplementary Fig. 9). For most α-Syn variants, the average condensate diameter decreased from ∼7 μm (0.5 μM α-Syn) to ∼5 μm (5 μM α-Syn)—implying a condensate volume reduction by a factor of ∼3 immediately after their formation (observed within 5 min). The inability of the Core variant (but not the ΔC1-125) to have this effect on DDX4N1 condensates suggested that the N-terminal region could be important for the formation of the Pickering clusters. We next hypothesized that clustering of negatively charged α-Syn on DDX4N1 condensate surfaces could prevent partitioning of nucleic acids within condensates by electrostatic repulsion, which could have crucial physiological significance. Using confocal microscopy, we measured the kinetics of partitioning of Alexa488-ssDNA into unlabeled DDX4N1 condensates in the absence/presence of 1 and 5 μM WT α-Syn (as Pickering clusters). In line with our hypothesis, we found a systematic decrease in ssDNA partitioning as a function of α-Syn concentration (Fig. 5d). The rate of partitioning decreased considerably, from 2.0 to 0.1 relative fluorescence unit (RFU)/s, in the presence of increasing α-Syn concentrations. Interestingly, a parallel FRAP experiment revealed that α-Syn did not alter the translational dynamics of DDX4N1 molecules within condensates (Supplementary Fig. 10)— indicating that the reduced diffusion of ssDNA was solely due to changes on the surface. Identical experiments with truncated α-Syn variants (ΔC1-125 and ΔC1-110) showed that the decreased partitioning of ssDNA was primarily caused by electrostatic repulsion from the acidic C-terminal of α-Syn, as this effect was less pronounced (from 2 to 1 RFU/s) for ΔC1-125 and ΔC1-110 compared to WT (Fig. 5e, Supplementary Fig. 11).

## DISCUSSION

In this body of work, we quantitatively study the phase separation behaviour of α-Syn, and compare it among important pathological modifications linked to PD^57,58^. Our experiments consistently show that most of the tested familial single-point mutants, except for A53T, do not induce significant alterations in the phase separation behavior of α-Syn. Familial mutations have been shown to alter lipid membrane binding and amyloid aggregation of α-Syn by altering the charge, hydrophobicity, or flexibility of the N-terminal helical region^58^. They also influence cross-elongation of amyloid fibrils where the structure of the template fibril (e.g., a mutant) shapes the structure of the resulting fibril, leading WT monomers to adopt features and toxicity of the mutant fibril^58-60^. Despite these effects, we do not *a priori* expect significant effects of these mutations on phase separation. This is because in contrast to fibril formation, where well-defined molecular contacts are key drivers and stabilisers of the fibrils, a constantly re-arranging network of multiple weak interactions drives protein-demixing^61,62^. In this context, it is plausible that these single point mutations would not substantially modify the overall driving forces for condensate formation, as the effects of such mutations can be compensated by new interactions enabled by the substitution. On the other hand, terminal truncations/alterations of α-Syn show a significant impact, having an inverted NaCl dependence compared to full-length α-Syn^35^ (Fig. 1). Low saturation concentrations (∼10 μM) across varying NaCl concentrations suggest a high likelihood for these variants to undergo demixing even within cellular crowded milieus during overexpression, and faulty protein homeostasis machinery due to aging. It could be that the effect of the familial point mutations are more aggressive and distinguishable on a truncated α-Syn backbone. Such combinatorial effects of pathological alterations remains to be investigated in the context of α-Syn phase separation, which we aim to address in a future study. In addition to their altered thermodynamic stability, truncated α-Syn condensates also represent more resilient, gel-like assembly states—exhibiting higher viscosity and resistance against dissolution immediately after their formation (Fig. 2). A natural question that arises from these findings is how these varying material properties influence or relate to the formation of PD associated amyloid fibrils within condensates. With the assumption that dense phase aggregation has a considerable faster kinetics of fibril formation compared to the dilute phase, we employed a sonication-based approach to release amyloid fibrils from condensates at particular timepoints where the majority of the dilute phase have not yet aggregated (Fig. 3). This allowed us to quantify and compare the apparent fibril concentrations (seeding potential) primarily from the dense phase of α-Syn variants. We believe this can be translated to many other IDP condensates undergoing solidification. The data from such experiments reveal an interesting aspect of truncated α-Syn condensates—that gelation of these assemblies, even within seconds, can give rise to the formation of elongation competent amyloid fibrils. Spontaneous sol-gel phase transition and amyloid fibril formation of truncated α-Syn could be particularly important, given the abundance of C-terminally truncated α-Syn aggregates in Lewy bodies present in PD affected individuals^63^. Our findings also demonstrate that despite significant sequence variations and differences in phase separating conditions, α-Syn consistently forms amyloid fibrils within condensates (Fig. 3)—suggesting that very different types of α-Syn condensates are likely similar in their nature to undergo amyloid aggregation upon gelation, provided that the amyloid-forming region (residues 60–100) is present in the protein sequence. Nevertheless, gelation of condensates into non-fibrillar aggregates could still be detrimental as accumulation of such condensates could stall/block protein degradation machineries, resulting in severe pathological consequences. In agreement with this line of thought, we find that irrespective of the differences in their dominant aggregation mechanism (across variants), most α-Syn amyloid fibril variants formed in the presence of condensates show considerable toxicity to human neuronal cell lines.

Biological condensates are multi-component and often enriched in nucleic acids for transcriptional and translational regulation^64,65^. Although in cell phase separation driven by intrinsic (as opposed to overexpressed) α-Syn (as a scaffold molecule) remains to be demonstrated, emerging evidence suggest that α-Syn can efficiently partition into other neuronal protein condensates (e.g., tau, synapsin and VAMP2) as a ‘client’ molecule^51-53^. However, the effects of α-Syn on nucleic acid enriched condensates/coacervates remain underexplored. We find that for both RP3/ssDNA coacervates^38^ and DDX4N1 model condensates^38,40^, the most prominent effect of α-Syn was imparted on nucleic acids (in our case: ssDNA)—modulated by the acidic C-terminal tail of α-Syn (Fig. 4-5). In neurons, the dissolution of phase-separated assemblies is equally (if not more) critical compared to their formation. In fact, the ability to form and dissolve repeatedly is a key advantage of a cellular condensate over other supramolecular assembly states. For many condensate systems, this dynamic cycling is mediated by enzymes (kinases) and ATP^66^. Here, we show another regulatory mechanism that might exist for many neuronal coacervates, where α-Syn (or α-Syn like proteins) can compete with ssDNA (or a negatively charged biomolecule) for RP3 peptide (or a positively charged protein), dissolving the coacervates by shifting the electrostatic balance of the system. On the other hand, α-Syn can also form Pickering clusters on the surface of DDX4N1 condensates. This phenomenon leads to significant changes in condensate size distribution, even at sub-stoichiometric ratios where α-Syn is 70 times less concentrated compared to DDX4N1. A similar effect was previously shown for TDP43 condensates where α-Syn emulsification resulted in TDP43 aggregation^56^. Here, we find that these Pickering clusters control condensate size and slow down ssDNA partitioning within DDX4N1 condensates, which might be implicated in the regulation and stabilization of DDX4N1 condensates against nucleic acid-induced dissolution. These insights demonstrate the multi-faceted nature of a single IDP on the condensation landscape of a living cell— both enabling and preventing dissolution of phase separated states depending on their molecular profile. Finally, this body of work validates the robustness of our in-house methods (Capflex^38^ and TDIPS^39^) to quantitatively study phase separation by running experimental campaigns to screen targeted mutational libraries, and understanding sequence-dependent effects.

## ACKNOWLEDGEMENTS

We thank Prof. Samir K. Maji (Department of Biosciences and Bioengineering, IIT-Bombay) for ΔC1-110 and Core α-Syn proteins and plasmids. We thank Rasmus K. Norrild for crucial insights leading to the discovery of the α-Syn Pickering clusters. DTU bioimaging core at DTU bioengineering is acknowledged for confocal/fluorescence imaging and FRAP experiments. Funding from Novo Nordisk Foundation (grant NNF19OC0055625) for the infrastructure “Imaging microbial language in biocontrol (IMLiB)” is acknowledged. DTU nanolabs is acknowledged for TEM imaging. A.K.B. would like to acknowledge funding through an ERC CoG (101088163 EMMA) for funding. S.R. would like to acknowledge Horizon MSCA individual postdoctoral fellowship (Grant number 110361) for funding. C.C. would like to acknowledge an Erasmus Masters scholarship. A.K.B. and A.F. would like to acknowledge the Michael J Fox foundation for funding (MJFF-021293) for funding. A.K. would like to acknowledge Horizon MSCA individual postdoctoral fellowship (Grant number 101106115XX) for funding. S.H. would like to acknowledge a Lundbeck foundation postdoctoral fellowship (Grant number R449-2023-1527) for funding. C.G. would like to acknowledge an Independent Research Fund Denmark DFF-Research Project 1 (Grant number: 3103-00220B) for funding. The funders had no role in study design, data collection and analysis, decision to publish or preparation of the manuscript.

## AUTHOR CONTRIBUTIONS

S.R., C.C, A.F., A.K., K.H.S., S.H., K.M., designed and performed experiments and analyzed data. L.K.K. expressed and purified α-Syn proteins. A.F. expressed and purified C_rev_ α-Syn. S.H., C.C., and S.R. purified DDX4N1. K.H.S., K.M., and C.G. conceptualized and performed all in cell experiments. A.K.B. acquired funding, conceived and supervised the study, designed experiments and analyzed data. S.R., C.C., K.H.S., and A.K.B. wrote the manuscript. S.R. prepared all schematics and illustrations. All authors commented on the manuscript and approved it.

## COMPETING INTERESTS

The authors declare no competing interests.

## Notes

### Competing Interest Statement

The authors have declared no competing interest.

