## Supplementary material for "Divergent effects of pathological α-synuclein truncations and mutations on phase separation": Supplemetary Information

### MATERIALS AND METHODS

**Materials:** WT and variant (A30P, H50Q, G51D, A53T, ΔC1-125, 140C) human α-Syn genetic sequences were encoded on a pT7-7 bacterial plasmid under isopropyl β-D-1 thiogalactopyranoside (IPTG) inducible promoter sequence (Genscript, USA). Purified ΔC1-110 and Core α-Syn variants were kind gifts from Prof. Samir K. Maji (IIT-Bombay, India). The $C_{rev}$ gene construct (D115, D121, E126, E131, and E137, all substituted to K) was purchased from Twist Bioscience (USA) on a pET-29b(+) plasmid with a stop codon placed before the His-tag at the end of the protein sequence. DDX4N1 CtoA gene was encoded as a GST fusion protein on a pET29a(+) bacterial plasmid (Genscript, USA). DDX4N1-YFP was encoded on pET30M-2 plasmid and was a kind gift from Dr. Tim Nott (Oxford University, UK). HPLC-purified $RP_3$ peptide (solid-phase synthesis) was procured from Bachem (Switzerland), and Schafer-N (Denmark). HPLC purified unlabeled, and Alexa488 labeled ssDNA was procured from TAG Copenhagen (Denmark). LB broth, ampicillin, kanamycin, and IPTG for protein production was procured from VWR (Denmark). All other relevant reagents, salts and buffer components for experiments were purchased from Sigma (USA) and VWR (Denmark), unless otherwise specified. Formvar coated electron microscopy grids were purchased from Sigma (USA). Protein, peptide and nucleic acid sequences are provided at the end of the method section.

#### Methods:

#### 1. Protein production:

**Expression and purification of α-Syn variants:** WT, A30P, H50Q, G51D, A53T, and ΔC1-125 α-Syn were produced and purified following identical protocols[1]. 100 mg/L ampicillin (kanamycin for familial point mutants) was used as a bacterial selection marker for protein production. After transformation with the desired plasmid, a primary *E. coli* BL21 (DE3) culture was grown for 10-12h at 37°C. The primary culture (10 ml) was then inoculated into 1 L LB and grown under a shaking speed of 160 rpm, till the $OD_{600}$ reached 0.8. Protein expression was induced by adding 1 mM IPTG and the culture was grown for 4 more hours. The cells were harvested by centrifugation (7000 xg, 20 min, 4°C) and the pellet was stored at -20°C until further use. To purify the protein, the cell pellet was dissolved in 20 ml, 10 mM Tris-HCl, 1mM ethylenediamine tetra acetic acid (EDTA), pH 8.0 with 1mM phenylmethylsulfonyl fluoride (PMSF) and sonicated on ice (10s on, 30s off, 12 cycles at 40% amplitude). Next, 1μL Benzonase was added to the cell lysate to precipitate DNA followed by centrifugation at 20000 xg for 30 min at 4°C. The supernatant was collected and heated at 80°C on a water-bath for 20 min. After cooling down the solution back to room temperature, a second centrifugation step (20000 xg for 20 min at 4°C) was performed to precipitate heat-sensitive proteins, while α-Syn remained in the supernatant. The supernatant was collected, and 4 ml saturated $(NH_4)_2SO_4$ was added for 1 ml supernatant to salt out α-Syn. Salted out α-Syn was obtained in the pellet after centrifuging the solution at 20000 xg for 20 min at 4°C. The pellet was dissolved in 7 ml of 25 mM Tris-HCl, pH 7.7, and 1 mM dithiothreitol (DTT) was added to the solution. Subsequently, the solution was dialyzed against the same buffer for 18h at 4°C to remove $(NH_4)_2SO_4$. The tank was replenished with fresh buffer after 12h. The α-Syn solution was then subjected to anion exchange column (AEC) (HiTrap Q Hp 5 ml, GE healthcare, USA) followed by size exclusion chromatography (SEC) (HiLoad 16/600 Superdex 200 pg. column). The purified protein was eluted in 10 mM of sodium phosphate ($NaH_2PO_4.H_20+Na_2HPO_4.2H_20$) buffer (pH 7.4). α-Syn concentrations were measured using a NanoDrop Lite (Thermo Scientific, USA), and a UV spectrophotometer, Labbot (Labbot, Sweden) by measuring the absorption at 280 nm. The theoretical molar extinction coefficient of full-length (5960 $M^{-1}cm^{-1}$) and ΔC1-125 α-Syn (2980 $M^{-1}cm^{-1}$) was predicted by ProtParam (Expasy, Switzerland). The protein purity was checked using SDS-PAGE. To note, 140C-α-Syn was expressed and purified using an identical protocol—the only difference being addition of 1 mM DTT and 1 mM EDTA in all buffers to prevent intermolecular cysteine-disulfide linkages[2].

**Expression and purification of C-terminal charge reversed ($C_{rev}$) α-Syn:** $C_{rev}$-α-Syn was purified in a small-scale setup, as previously established in our laboratory[1]. Briefly, the cell pellet obtained from 50 ml of culture was resuspended in 4 ml Tris buffer (10 mM, 1mM EDTA, pH 8) and 2 mM PMSF. The cells were sonicated on ice (10s on, 30s off, 12 cycles at 40% amplitude) and centrifuged at 20000 xg for 20 min, 4°C. The supernatant was collected and heated to 85°C for 20 min, followed by centrifugation (20000 xg for 20 min, 4°C). Next, DNA was precipitated by adding 5% v/v glacial acetic acid and 1 mg/ml streptomycin sulfate, followed by centrifugation (20000 xg for 20 min, 4°C). The supernatant was transferred into another tube and $C_{rev}$ α-Syn was salted out with saturated $(NH_4)_2SO_4$ (final concertation of 2.5 M). The pellet was gently washed 2 times and dissolved in 10 ml Tris buffer (10 mM, pH 8.0). The protein was loaded on a gravity column with cation exchange resin (NuviaTM S, BIO-RAD, USA). The column was washed with 4 ml of 150 mM NaCl and the protein was eluted with 3 ml of 250 mM NaCl (10 mM Tris pH 8.0). The buffer was exchanged with an illustra NAP-5 column (GE Healthcare, USA) in 10 mM of sodium phosphate ($NaH_2PO_4.H_20+Na_2HPO_4.2H_20$) buffer at the desired pH, before use. Protein concentration was measured by NanoDrop Lite (Thermo Scientific, USA) using 5960 $M^{-1}.cm^{-1}$ as the molar extinction coefficient for $C_{rev}$ α-Syn. The protein purity was checked using SDS-PAGE.

**DDX4N1 expression and purification:** A variant of DDX4N1 was used in our experiments where all cysteine residues were mutated to alanines (CtoA)[1]. This avoids disulphide bridge formation and also simplifies the protein purification protocol, having little or no effect on LLPS. We refer to this variant as only 'DDX4N1' for better readability. DDX4N1 was tagged with a His_tag fused with a thioredoxin-tag (eMM9) at the C-terminal that facilitates protein expression and thermodynamic stability. The His_tag-eMM9 was followed by codon-optimized DDX4N1 CtoA linked by a 10-residue GS-linker and a TEV-cleavage site (ENLYFQ/G). For DDX4N1-YFP, the protein was produced as His_tag-GST-TEV_site-DDX4-YFP. Successful clones were selected from a transformation plate (kanamycin as selection marker) and both the proteins were produced and purified following similar, previously described established protocols[1,2]. Briefly, for DDX4N1, 10 ml overnight grown culture was (50 mg/l kanamycin) was used to inoculate 1l AB-LB. The AB-LB-medium contained 5 g/l NaCl, 40 mM sodium/potassium phosphate (K/NaPi) buffer, pH 7.0, 15 mM $(NH_4)_2SO_4$, 50 mM NaCl, 2 mM $MgCl_2$, 0.1 mM $CaCl_2$, 3 nM $FeCl_3$, and 50 mg/l kanamycin. The culture was grown at 37°C till $OD_{600}$ reached 0.8 and protein production was induced with 1 mM IPTG, before incubating overnight at 20°C. Cells were harvested by centrifugation at 7000 xg for 20 min at 4°C. 30 ml lysis buffer (50 mM sodium phosphate buffer, pH 6.5, 500 mM NaCl) was used to resuspend the cell pellet. 8 µl Benzonase was added to the solution and the resuspended cells were lysed on ice with the help of a probe sonicator (40% amplitude, 30s on, 30s off, total time of 30 min). The cell lysate was centrifuged at 20000 xg for 20 min at 4°C and the supernatant was collected and heated at 80°C for 15 min. The solution was centrifuged at 4500 xg for 30 min at 4°C to remove heat-induced protein aggregates and the supernatant was passed through a 0.22 µm filter and collected. 10 mM imidazole was added and the solution was loaded onto 5 ml pre-equilibrated Ni-NTA resin (Thermo Scientific) on a gravity column. The column was washed with 10 column volumes of 50 mM sodium phosphate, pH 6.5, 500 mM NaCl, 20 mM imidazole, followed by 10 column volumes denaturing buffer (50 mM sodium phosphate, pH 6.5, 500 mM NaCl, 20 mM imidazole, and 3M GdnHCl). Next, GdnHCl was removed by washing the column with 10 column volumes of the non-denaturing buffer. A protease solution (9.5 ml lysis buffer, 2 mM TCEP, 500 µl His_tag-TEV protease (Genscript) was added to the column to release DDX4N1 overnight at room temperature with gentle mixing. TEV protease remained bound, and the eluted DDX4N1 was concentrated using a 10 kDa centrifugal filter (Amicon ultra, Merck, Germany) to 2 ml. The concentrated DDX4N1 was SEC purified on a HiLoad Superdex 75 16/600 column. Pooled fractions were flash frozen in liquid nitrogen before storing at -80°C. The protein purity was checked at different stages of purification using SDS-PAGE. The DDX4N1 concentrations were measured using by NanoDrop Lite (Thermo Scientific, USA) using molar extinction coefficients of 29350 $M^{-1}.cm^{-1}$ determined from ProtParam (Expasy, Switzerland).

**Fluorophore labeling of proteins:** In this work, we have used Alexa488-140C-α-Syn, N-terminally Alexa488 labeled WT and ΔC1-125 α-Syn, Alexa488-ssDNA, and DDX4N1-YFP as fluorescent reporter molecules. Alexa488-ssDNA was purchased from a commercial supplier as mentioned in the materials section. DDX4N1-YFP was recombinantly produced in *E.coli* as a fusion protein[2]. For Alexa488-140C-α-Syn production, 13.7 mg/ml 140C-α-Syn was injected into a Superdex 200 increase, 10/300 GL column to remove DTT and EDTA. The fraction containing 140C-α-Syn was pooled at a concentration of 5.8 mg/ml. Alexa488 C-5 maleimide was dissolved in DMSO (1 mg in 200 μl) which corresponds to a concentration of ~7 mM. The labeling reaction was performed at 25°C for 1h, with 360 μM 140C-α-Syn and 10 fold excess (3.6 mM) dye concentration. The total reaction volume was 500 μl. After conjugation, the free dye was removed using Superdex 200 increase, 10/300 GL column and the concentration of the labeled protein was calculated according to absorbance values at 275 nm (140C-α-Syn) and 488 nm (Alexa488 C5 maleimide) with the help of NanoDrop Lite (Thermo Scientific, USA), and using 5960 $M^{-1}.cm^{-1}$ as the molar extinction coefficient of the protein. N-terminal Alexa488-WT/ΔC1-125 α-Syn were produced using labeling kits provided by the dye supplier: FIDAbiosystems ApS (Denmark). Here, the Alexa488-NHS labeling reaction was performed in sodium phosphate buffer (pH 7) instead of bicarbonate buffer (pH ~9, used to label the lysine side chain amines) to specifically target the primary amine group at the N-terminal of the protein.

**2. In vitro phase separation assays:**

**TDIPS experiments with α-Syn**: Taylor dispersion induced phase separation (TDIPS) is an in-house developed, high-throughput capable phase separation screening platform based on the FIDA1 instrument (FIDAbiosystems ApS, Denmark)[3]. This method was used to chart the orthogonal phase diagrams (as a function of NaCl and protein concentration) of α-Syn variants. A small plug (tens of nl) of α-Syn (different concentrations, with 25% (w/v) PEG-8000, in 20 mM sodium phosphate buffer, pH 7.4) was injected into the FIDA1 capillary (75 μm diameter, 100 cm length) and sandwiched between the same buffer (20 mM sodium phosphate buffer, pH 7.4, 25% (w/v) PEG-8000) containing different concentrations (0-250 mM) of NaCl to induce phase separation. The experimental parameters are described in the following table. The experiments were conducted at room temperature (25°C).

**Supplementary Table 1: TDIPS parameters for α-Syn phase diagrams:**

| Tray | Vial | Pressure (mbar) | Time (s) | Outlet | Measure | Comment |
|---|---|---|---|---|---|---|
| 2 | 1 | 3500 | 360 | Variable | no | 1 M NaOH wash for cleaning the capillary. |
| 2 | 2 | 3500 | 180 | Variable | no | MQ water wash to remove NaOH. |
| 2 | 3 | 3500 | 30 | Variable | no | 20 mM sodium phosphate buffer, pH 7.4 to equilibrate the capillary. |
| 1 | Analyte (NaCl as variable) | 3500 | 90 | Variable | no | 20 mM sodium phosphate buffer, pH 7.4, 25% (w/v) PEG-8000, with appropriate salt concentration as the experimental condition. |
| 1 | Indicator (Protein) | 900 | 10 | Variable | no | 200 μM α-Syn+20% (w/v) PEG-8000 in 20 mM sodium phosphate buffer, pH 7.4. Each sample was spiked with 20 nM Alexa488140C-α- |

| | | | | | | |
|---|---|---|---|---|---|---|
| | | | | | | Syn as fluorescent reporter molecule. |
| 1 | Analyte (NaCl as variable) | 2800 | 600 | Variable | yes | 20 mM sodium phosphate buffer, pH 7.4, 25% (w/v) PEG-8000, with appropriate salt concentration as the experimental condition. |

In the TDIPS traces, condensates (as fluorescent spikes) populated the trace at a time window from 3 to 4 minutes (Supplementary Fig.1). The extent of phase separation was calculated by multiplying the number of detected spikes (n) to the average intensity of the spikes ($I_{avg}$, reflecting the average size/volume of condensates). A threshold of 15% above the baseline signal was selected to find spikes in the TDIPS traces. The spike number and average spike intensity was calculated using a bult-in peak analyzer tool in Origin Pro software (Origin Labs, USA).

**TDIPS experiments with $RP_3$/ssDNA**: The $RP_3$/ssDNA TDIPS experiments were performed using a triple-plug strategy, where a small plug of $RP_3$ was placed between two identical ssDNA containing plugs (same volume as the $RP_3$ plug). This triple-plug was sandwiched between a buffer solution of identical ionic strength and pH of the plugs. We performed all experiments with 10 mM Tris-HCl buffer, pH 8.0, in presence of 150 mM NaCl. α-Syn variants were titrated in the ssDNA plugs. The experimental parameters are described in the following table. The experiments were conducted at 20°C.

**Supplementary Table 2: TDIPS parameters for $RP_3$/ssDNA coacervation:**

| Tray | Vial | Pressure (mbar) | Time (s) | Outlet | Measure | Comment |
|---|---|---|---|---|---|---|
| 2 | 1 | 3500 | 45 | Variable | no | 1 M NaOH wash for cleaning the capillary. |
| 2 | 2 | 3500 | 45 | Variable | no | MQ water wash to remove NaOH. |

| | | | | | | |
|---|---|---|---|---|---|---|
| 2 | 3 | 3500 | 45 | Variable | no | 10 mM Tris-HCl buffer, pH 8.0, 150 mM NaCl to equilibrate the capillary. |
| 1 | Analyte | 50 | 15 | Variable | no | 10 μM ssDNA spiked with 240 nM Alexa488-ssDNA in 10 mM Tris-HCl buffer, pH 8.0, 150 mM NaCl. This plug had different concentrations of α-Syn variants as titrants. |
| 1 | Indicator | 50 | 15 | Variable | no | 1.5 mM $RP_3$ in 10 mM Tris-HCl buffer, pH 8.0, 150 mM NaCl. |
| 1 | Analyte | 50 | 15 | Variable | no | 10 μM ssDNA spiked with 240 nM Alexa488-ssDNA in 10 mM Tris-HCl buffer, pH 8.0, 150 mM NaCl. This plug had different concentrations of α-Syn variants as titrants. |

| | | | | | | |
|---|---|---|---|---|---|---|
| 2 | 3 | 1000 | 90 | Variable | yes | 10 mM Tris-HCl buffer, pH 8.0, 150 mM NaCl to mobilize the triple-plug to the detector. |

In the absence of and at low stoichiometric ratios of α-Syn, $RP_3$ and ssDNA underwent electrostatic coacervation, giving rise to a splitting of an otherwise Gaussian profile of the detected ssDNA signal. The splitting of the Gaussian profile stems from mass transfer due to phase separation because condensates that form in front and at the back of the plug have essentially zero diffusivity compared to the monomeric peptide and DNA molecules[3]. The maximum peak intensity without phase separation was selected as a reference point and the area underneath the peak(s) on either side ($A_1$ and $A_2$, Supplementary Fig.7) were integrated using a bult-in peak analyzer tool in Origin Pro software (Origin Labs, USA). The ratio of the areas under the leading and the trailing peaks ($A_1/A_2$) was used as an indication of phase separation as a function of α-Syn concentrations relative to $RP_3$.

**Capflex experiments with α-Syn**: Capillary flow experiments (Capflex[2]) is another in-house developed phase separation platform based on the FIDA1 instrument (FIDAbiosystems ApS, Denmark) where we can measure dilute phase protein concentrations ($C_{dil}$) of a phase separating system within a capillary, without physically separating the dense and the dilute phases. Here, we quantify the relative decrease of the baseline fluorescence signal upon phase separation, compared to a non-phase separated sample of identical concentration of protein (Supplementary Fig.1). Presence of condensates can also be confirmed by emergence of fluorescence spikes—each one corresponding to an individual condensate passing the detector, and the spike intensity scales with the size/volume of the condensate. A calibration curve is generated from the baseline fluorescence values of known protein concentrations (without phase separation), which is used to determine the $C_{dil}$ of a phase separating solution. Using Capflex, we measured the $C_{dil}$ of different α-Syn variants under a range of solution conditions. For the experiments described in Fig.1, and Supplementary Fig.2, we induced phase separation in presence of 20% (w/v) PEG-8000, 20 mM sodium phosphate buffer, at different NaCl concentrations and pH. We quantified $C_{dil}$ using 20 nM Alexa488-140C-α-Syn as a fluorescent reporter. Unlike TDIPS where phase separation occurs within the capillary, in Capflex, we induced phase separation by pre-mixing the necessary components in 1 ml glass vials (FIDAbiosystems ApS, Denmark) and/or 96 well-plates which are compatible with the FIDA1 instrument and immediately injected the samples within the capillary for detection. The parameters used for these experiments are provided in the following table. All experiments were performed at 25°C.

**Supplementary Table 3: Capflex parameters for α-Syn phase separation:**

| Tray | Vial | Pressure (mbar) | Time (s) | Outlet | Measure | Comment |
|---|---|---|---|---|---|---|
| 2 | 1 | 3500 | 350 | Variable | no | 1 M NaOH wash for cleaning the capillary. |
| 2 | 2 | 3500 | 120 | Variable | no | MQ water wash to remove NaOH. |
| 2 | 3 | 3500 | 60 | Variable | no | 20 mM sodium phosphate buffer equilibration with appropriate NaCl concentration and pH as the analyte (phase separated solution). |
| 1 | Analyte (Phase separated solution) | 2800 | 600 | Variable | yes | α-Syn variants with 20% (w/v) PEG-8000 at different NaCl concentrations and pH. Each sample was spiked with either 20 nM Alexa488-140C-α-Syn unless otherwise mentioned. |

We analyzed all datasets using Origin Pro software (Origin Labs, USA).

**Capflex experiments with DDX4N1**: For DDX4N1 phase separation, the capillary was coated with 0.1% (v/v) Tween-20 before the beginning of every run to prevent the condensates from sticking to the walls. The experiments were carried out at room temperature (25°C), in 10 mM sodium phosphate buffer, pH 6.5, and 50 mM NaCl. The total DDX4N1 concentration was 70 µM, with 600 nM DDX4N1-YFP as a fluorescent reporter. The $C_{dil}$ of DDX4N1 was measured as a function of α-Syn concentrations (Fig.5). The experimental parameters are provided in the following table.

**Supplementary Table 4: Capflex parameters for DDX4N1 phase separation:**

| Tray | Vial | Pressure (mbar) | Time (s) | Outlet | Measure | Comment |
|---|---|---|---|---|---|---|
| 2 | 1 | 3500 | 200 | Variable | no | 1 M NaOH wash for cleaning the capillary. |
| 2 | 2 | 3500 | 60 | Variable | no | MQ water wash to remove NaOH. |
| 2 | 3 | 3500 | 40 | Variable | no | 0.1% (v/v) Tween-20 coating to prevent condensate sticking on the capillary wall. |
| 2 | 4 | 3500 | 40 | Variable | no | 10 mM sodium phosphate buffer, pH 6.5, 50 mM NaCl to equilibrate the capillary. |
| 1 | Analyte (Phase separated solution) | 500 | 250 | Variable | yes | 70 µM DDX4N1 phase separated solution with different concentrations of |

WT α-Syn. Each sample was spiked with 600 nM DDX4N1-YFP.

---

**Dilute phase concentration measurements using centrifugation:** To verify whether the Alexa488-140C-α-Syn could be used universally across the protein variants (in Capflex) for measuring $C_{dil}$ of phase separation, we quantified $C_{dil}$ using centrifugation assays. Phase separation of α-Syn variants were induced at 200 μM total protein concentration, in presence of 20% (w/v) PEG-8000, in 20 mM sodium phosphate buffer, at different NaCl concentrations, and at 25°C. α-Syn condensates are particularly difficult to spin down using centrifugation due to the presence of high PEG concentrations in the system. The centrifugal acceleration required to spin down spherical protein condensates by a distance 'h' is given by the following equation: $n.g = \frac{6\pi\eta r.h}{t.(\rho_d - \rho_s)V.}$. Here, the product of 'n' and 'g (acceleration due to gravity)' denotes the centrifugal RCF value; '$\rho_d$' is the density of protein inside the condensate; '$\rho_s$' is the density of the solvent; 'V' is the volume of the condensate; 'r' is the radius of the condensate, 't' is time, and 'η' is the viscosity of the solvent[1]. We assumed that the density of condensates to be: $\rho_d$~1.1 g/cm$^3$ (at 25°C) and $\rho_s$=1.01 g/cm$^3$ (at 25°C). Therefore, to move even the smaller α-Syn condensates having an arbitrary radius of 0.3 μm by 1 cm in a 20% (w/v) PEG-8000 solution (η=20 mPa.s) within 2h, required an RCF value of ~15500 xg. We used 16000 xg RCF for 2h to ensure most α-Syn condensates are separated in pellets, and measured the $C_{dil}$ using a NanoDrop Lite (Thermo Scientific, USA). Importantly, thermodynamics of α-Syn phase separation is less sensitive towards temperature change (upto 65°C) compared to other canonical systems[1]. Therefore, we could perform centrifugation at 4°C without risking change in the $C_{dil}$ in response to temperature change. Identical centrifugation parameters were also used to isolate the dense phase for amyloid fibril seeding experiments described in Fig.3 and Supplementary Fig.5.

**Static light scattering and fluorescence spectroscopy**: SLS and spectroscopy experiments (Fig.2 and Supplementary Fig.4, respectively) were carried out using a thermally controlled, multichannel spectrophotometer—Labbot (Labbot, Sweden). For SLS measurements, 200 μM α-Syn variants were phase separated in presence of 20% (w/v) PEG-8000, in 20 mM sodium phosphate buffer, pH 7.4, at different NaCl concentrations (optimal for phase separation for each variant). Total volumes of the reaction mixtures were ~70 μl for each variant. The samples were dispensed in a 3x3 mm light path quartz glass cuvette (Hellma Analytics, Germany) and measured for 40 minutes at a constant temperature of 25°C. Immediately after loading the samples, 1 M Urea (prepared in 20 mM sodium phosphate buffer, pH 7.4) was added from an 8 M stock solution to induce condensate dissolution. The dilution of the protein and PEG concentrations by the addition of the urea solution does not place the composition of the sample outside of the two-phase region of the phase diagram. We measured SLS at 90° with a 636 nm laser with a 20 s equilibration time and 30 s interval time between each measurement, over 40 minutes. The sample was mixed every few minutes using a micropipette to prevent sedimentation of the condensates. For the experiments described in Supplementary Fig.4c-d, 200 µM monomeric α-Syn solution (no PEG) in 20 mM sodium phosphate buffer, pH 7.4, with 200 µM ThT was sonicated in 1.5 µl microcentrifuge tubes (Eppendorf) using a no-contact ultrasound probe at 80% amplitude, 1 sec on/3 sec off, for a total duration of 10 minutes (on ice, 4°C). 60 µl of this sonicated sample was dispensed in a 3x3 mm light path quartz glass cuvette (Hellma Analytics, Germany) and the emission spectra (440-600 nm) was measured with an excitation wavelength of 405 nm. Subsequently, 1 µM of pre-formed amyloid fibrils were added to the same sample and the emission spectra was measured again. This was to confirm fibril binding ability of ThT was retained after sonication.

**3. Thioflavin-T aggregation assays:**

**ThT aggregation kinetics of α-Syn condensate solutions:** ThT kinetics experiments described in Fig.3 were performed with 200 µM α-Syn variants, 20% (w/v) PEG-8000, in 20 mM sodium phosphate buffer, pH 7.4, and at different NaCl concentrations. The samples were dispensed in low-binding 96 well plates (Thermo Scientific, USA) and measurements were conducted under quiescent conditions, at 25°C, with the help of a fluorescence microplate reader: FLUOstar Omega (BMG Labtech, Germany). The surrounding wells were filled with MQ water to prevent evaporation. Each sample contained monomer equivalent concentration of ThT (i.e., 200 µM). ThT fluorescence over time was monitored every 15 minutes for ~7 days, by exciting the samples at 440 nm and recording the emission at 482 nm. Notably, 0.05% (w/v) sodium azide was added to all samples to prevent bacterial/fungal contamination over long incubation periods.

**Estimation of templating/elongation competent fibril concentration in α-Syn condensate solutions:** 150 µl of 200 µM α-Syn variants, 20% (w/v) PEG-8000, in 20 mM sodium phosphate buffer, pH 7.4, and at different NaCl concentrations (optimal for their respective phase separation) were prepared. The solutions were dispensed in low-binding 96 well plates (Thermo Scientific,

USA) and ThT fluorescence was measured as described above with a FLUOstar Omega (BMG Labtech, Germany). After 48h of incubation (with measurements every 15 min), WT, A30P, H50Q, G51D, A53T α-Syn condensate solutions were shifted to 1.5 ml microcentrifuge tubes and sonicated using a no-contact ultrasound probe (VialTweeter) at 80% amplitude, 1 sec on/3 sec off, for a total duration of 10 minutes (on ice, 4°C). The sonicated samples were dispensed back into the same wells in the 96 well plates, and ThT fluorescence measurements were resumed under quiescent conditions (25°C). For Core and ΔC1-125 α-Syn, the condensate solutions were sonicated at 0h (few minutes after their formation) with the same setting as mentioned above. ThT fluorescence values for 4-5 hours post sonication was fit to a linear equation and the slopes (elongation rates) of the fit lines were calculated. In parallel, calibration curves were generated for each α-Syn variant with known (0.5, 1 and 5 μM) monomer equivalent amyloid fibril concentrations. The calibration experiments were performed with 30 μM monomeric α-Syn variants, 15% (w/v) PEG-8000, in 20 mM sodium phosphate buffer, pH 7.4, and at identical NaCl concentrations as the phase separated samples (0 mM for truncated and 250 mM for full-length variants). A lower monomer and PEG concentration in the calibration experiments was to ensure no α-Syn variant was phase separated to a significant degree, and the majority of the dilute phase monomers were available for fibril elongation. The viscosity difference arising from 15% (w/v) PEG-8000 in the calibration experiment, versus 20% (w/v) PEG-8000 in tests could not be avoided since increasing the PEG concentration in calibration experiments could lead to extensive phase separation of the truncated variants, potentially lowering the $C_{dil}$ significantly (therefore, changing the elongation rates). This was not a major concern since the results were focused on comparing apparent seeding potential of condensates formed by different α-Syn variants, and does not reflect an absolute quantification of the amyloid fibril concentrations. The amyloid fibrils used as elongation competent seeds in the calibration experiments were prepared by constantly shaking (700 rpm, with a glass bead, for 7 days at 37°C) 50 μM monomeric α-Syn variants, in 20 mM sodium phosphate buffer, pH 7.4, and at identical NaCl concentrations as the phase separated samples. These fibrils were sonicated by following the same protocol discussed above for condensate solutions. The elongation rate constant ($k_+$) for each α-Syn fibril variant was calculated by measuring the elongation rates ($\frac{d[ThT\ fluorescence]}{dt}$) at different, known fibril concentrations [F] by the following equation: $\frac{d[ThT\ fluorescence]}{dt} = k_+[F]$. From this, the approximate monomer equivalent fibril concentrations in condensate (unknown) samples were calculated.

In a different approach (Fig.3e), we centrifuged condensate solutions (prepared under identical solution conditions as described above) of α-Syn variants in 1.5 μl microcentrifuge tubes at 48h (full-length) and 0h (truncated). We assumed that the lower limit of condensate radius could be 0.3 μm and therefore, used a centrifugal acceleration of 16000 xg for 2h, at 4°C to collect most of the dense phase. The supernatant containing the dilute phase monomers and PEG was discarded and a fresh monomeric (30 μM) α-Syn solution (identical buffer and NaCl concentration as condensate solutions) was added to each condensate pellet. Note that the monomeric solution did not contain PEG. Additionally, from our previous experiments, we know that PEG remains excluded from α-Syn condensates[1]. Therefore, at this point, the solutions contained very little PEG. The pellet was resuspended and subsequently sonicated (80% amplitude, 1 sec on/3 sec off, 10 minutes, 4°C) to disrupt the condensates, potentially releasing templating competent amyloid fibrils within them. 30 μM ThT was added and the sonicated solutions were dispensed in a low-binding 384 well plate (Thermo Scientific, USA). ThT fluorescence was measured with a FLUOstar Omega (BMG Labtech, Germany) for 30h under quiescent conditions, at 25°C. In parallel, calibration curves were generated with 0.1, 0.5 and 1 μM pre-made α-Syn fibril seeds and 30 μM monomeric proteins for each α-Syn variant (without PEG). The elongation rate constant ($k_+$) for each α-Syn fibril variant was calculated and the approximate monomer equivalent fibril concentrations (seeding potential) of condensates (unknown) were determined. This method had two advantages over the method where we measured seeding potential without isolating the condensates (Fig.3d), but also had one major disadvantage. First, the calibration curves could be generated without PEG, mitigating the risk of phase separation especially for the truncated variants. Second, the monomer concentration in the experiments were identical to the calibration curves. However, harvesting the dense phase entirely could be challenging in PEG solutions, especially for sub-micron/nanoscale condensates[1] which might contribute to a substantial fraction of the dense phase. This is likely why we found reduced apparent fibril concentrations for all α-Syn variants in this version of the assay, and a substantial variation between repeated measurements for G51D (Fig.3f).

**4. Microscopy:**

**Microscopic observation of phase separated condensates**: For α-Syn, condensate solutions containing 200 μM α-Syn variants, 20% (w/v) PEG-8000, in 20 mM sodium phosphate buffer were prepared at different pH and NaCl concentrations (optimal for phase separation). 100 nM Alexa488-140C-α-Syn was used as a fluorescent reporter molecule. For $RP_3$/ssDNA, phase separated coacervates were prepared by mixing 200 μM $RP_3$ with 20 μM ssDNA, and as a function of WT and

ΔC1-125 α-Syn. These experiments were performed in 10 mM Tris-HCl buffer, pH 8.0, in the presence of 150 mM NaCl. 100 nM Alexa488 ssDNA or N-terminally labeled Alexa488-WT and Alexa488-ΔC1-125 α-Syn were used as fluorescent reporters (as mentioned for different experiments). For DDX4N1, 70 µM protein was phase separated by rapidly diluting a 700 µM stock solution (at 500 mM NaCl) tenfold to 50 mM NaCl, in 10 mM sodium phosphate buffer, pH 6.5. 100 nM DDX4N1-YFP was used as a fluorescent reporter. To visualize α-Syn Pickering clusters on DDX4N1 condensates (Fig.5), 100 nM of Alexa488-140C-α-Syn was used as a fluorescent reporter. All experiments described above were visualized with an LMI-005-Confocal Microscope SP8 (Leica Microsystems, Germany). Images were acquired immediately after dispensing 5 µl phase separated solution to 15 µl PDMS wells on glass coverslips, with a 63X (oil immersion) objective at a resolution of either 512/512 or 1024/1024 pixels, and at 16 bit-depth. The excitation wavelength was set at 480 nm and images were captured with an emission range of 500-700 nm. The laser exposure was adjusted for individual sample so that maximum number of condensates/clusters could be detected. All experiments were performed at 25°C.

**Fluorescence recovery after photobleaching:** FRAP experiments with α-Syn and DDX4N1 condensates were performed according to previously established protocols[4,5]. Briefly, a bleaching radius of 1-3 µm was chosen depending on the α-Syn/DDX4N1 condensate size. The condensates were photobleached with a 488 nm laser at 100% power for 2s. The fluorescence recovery was recorded at a 371 ms frame rate for 20-90 s (post-bleach) and subsequently corrected for the effect of passive bleaching and the background fluorescence. The post-bleach fluorescence intensity values were normalized with respect to the pre-bleach fluorescence for individual condensates. All experiments were carried out at 25°C. The images were processed and analyzed using ImageJ (NIH, USA). 100 nM of Alexa488-140C-α-Syn and DDX4N1-YFP were used as a fluorescent reporter, for α-Syn and DDX4N1 condensates, respectively. All data were plotted and statistical significance was tested using Origin Pro (Origin Labs, USA).

**Condensate fusion analysis:** Fusion events of α-Syn and DDX4N1 condensates were recorded using a widefield Zeiss Axio vert A1 microscope (Zeiss, Germany) in bright-field mode, with a 40x objective lens. A resolution of 1626/1236 pixels with a frame rate of 50 ms were chosen. Images were acquired immediately after dispensing 5 µl phase separated solution to 15 µl PDMS wells on glass coverslips. DDX4N1 fusion events were obtained from phase separated samples at a concentration of 70 µM, in 10 mM sodium phosphate buffer, pH 6.5, 50 mM NaCl (in the absence and presence of 5 μM α-Syn). Fusion events of WT and ΔC1-110 α-Syn condensates were recorded in phase separated solutions containing 200 μM α-Syn variants, 20% (w/v) PEG-8000, in 20 mM sodium phosphate buffer, pH 7.4, at optimal NaCl concentrations. We noted that fusion events were rare in phase separated solutions and could only detect tens of events over 3 independent experiments. This was likely because of the high viscosity of the solution (slower diffusion) due to presence of 20% (w/v) PEG-8000. In general, timescales of fusion events of liquid-like condensates are used to determine the ratio between the viscosity (η) and the surface tension (γ) of the condensates—known as the inverse capillary velocity ($\nu = \eta/\gamma$)[6,7]. When two spherical condensates of similar sizes are in contact (as in our case), a hypothetical ellipse is drawn which has a major axis of $D_0$ ($D_0/2$, being the diameter of each condensate just before fusion, at time=0). Let the minor axis of this ellipse be $S_0$. During fusion, the aspect ratio of the ellipse, i.e., the value of $D_0/S_0$ exponentially decays with the relaxation time, reaching a value of 1 after complete relaxation into a larger condensate. Fitting this data to a mono-exponential decay function is used to calculate the characteristic time (τ, exponential constant). For each condensate pair undergoing fusion, the geometric mean diameter, or the lengthscale ($l_0$) of fusion (at time=0) is defined with the following equation: $l_0 = \sqrt{S_0(D_0 - S_0)}$. Plotting $l_0$ as a function of τ for a series of fusion events, and fitting the data with a linear equation is then used to calculate ν, because $\tau/l_0 = \eta/\gamma = \nu$. In our case, we used a simplified, comparative analysis where we considered only the change in $D_0$ as a function of relaxation time—the aspect ratio being approximately proportional to $D_0$, and the $l_0$ being proportional to $\sqrt{D_0}$. The distances are measured along the diameters of fusing condensates, i.e., from one end of a condensate to the other end of the other condensate (Supplementary Fig.3). End-to-end distances of detected fusion events as a function of relaxation time were fitted with a mono-exponential decay function: $D(t) = D_0 e^{-t/\tau_{app}} + C$, where D(t) corresponds to distance at time=t, $D_0$ corresponds to distance at time=0, $\tau_{app}$ is the apparent time constant of fusion events ($\tau_{app}=k\tau$, where k is a proportionality constant relating $D_0$ and the aspect ratio). C is the long-term asymptote, i.e., the value of D(t) when t approaches infinity. We then plotted $\tau_{app}$ as a function of $\sqrt{D_0}$ to calculate the apparent inverse capillary velocity ($\nu_{app}$) for WT and ΔC1-110 α-Syn condensates. The images were analyzed using ImageJ (NIH, USA) and Origin Pro (Origin Labs, USA).

**Nucleic acid partitioning within DDX4N1 condensates:** The PDMS wells on glass coverslips[1] were filled with 6 μl of unlabeled 70 μM phase separated DDX4N1 solution prepared in 10 mM phosphate buffer, pH 6.5, 50 mM NaCl. Subsequently, 1 μl of 500 nM Alexa488-ssDNA was dispensed to the well to monitor partitioning of labeled DNA within unlabeled DDX4N1 condensates (in the presence and absence of 5 μM α-Syn variants: WT, ΔC1-125, ΔC1-110). The samples were visualized with an LMI-005-Confocal Microscope SP8 (Leica Microsystems, Germany) with a laser power of 5%, gain of 800, PMT gain of 250, and a framerate of 1.2s/frame. The data were analyzed using ImageJ (Fiji, NIH, USA). The fluorescence intensity from each condensate was averaged and background corrected for N~50 individual condensates at each time point, and the values were plotted as a function of time (Fig.5, Supplementary Fig.11). The slopes were calculated using linear fitting of the data using Origin Pro (Origin Labs, USA).

**Transmission electron microscopy (TEM):** TEM imaging of condensate derived amyloid fibrils of α-Syn variants was performed following previously established protocols[1]. The samples were aged for 7 days at 25°C under optimal phase separating conditions. Before imaging, the samples were diluted to 50 μM total protein concentration (monomer equivalent) using 20 mM sodium phosphate buffer, pH 7.4, and 10 μl of these diluted solutions were drop casted onto formvar coated TEM grids (Sigma, USA). The grids were left for drying at room temperature for 10 min. The excess solution was blotted carefully using a Whatman filter paper without touching the grid. Next, 10 μl 1% (w/v) uranyl acetate (prepared in MQ water) was drop casted on the grids and the samples were negatively stained for 30s. Excess dye was blotted using filter paper and the grids were left for drying at room temperature for 20 min. The grids were subsequently imaged using a 200 kV Tecnai T20 G2 electron microscope (FEI, USA) at desired magnification (10000x). The images were captured using a TVIPS XF415 CMOS 4K camera and TVIPS EMplify v0.4.5software (50-100 ms exposure time depending on the sample).

**5. Cellular toxicity:**

**MTT cell viability assay on SH-SY5Y cells:** Condensate solutions containing 200 μM α-Syn variants, 20% (w/v) PEG-8000, in 20 mM sodium phosphate buffer (no azide), pH 7.4, and at different NaCl concentrations (optimal for phase separation) were prepared, and aged in 1.5 ml microcentrifuge tubes (Eppendorf) for 7 days at 25°C to allow sufficient protein aggregation. Notably, the full-length variants were sonicated at 48h and the truncated variants were sonicated at 0h, to allow propagation of the condensate derived amyloid fibrils primarily via elongation mechanisms. Sonication was performed following the same protocol as described before, in no contact mode. Undifferentiated SH-SY5Y cells were seeded at 25000 cells/well in a cell culture grade 96-well plate. The condensates derived amyloid fibrils/aggregates made from α-Syn variants and corresponding α-Syn variant monomers were diluted to 5 μM in cell media (DMEM GlutaMAX, high glucose with 10% (v/v) Foetal bovine serum (FBS), 1% (v/v) Penicillin/Streptomycin, and 1% (v/v) non-essential amino acids (NEAA). The condensate samples were sonicated again to fragment amyloid fibrils just before adding to the cells. After either 24h or 96h, the media was removed and thiazolyl blue tetrazolium bromide (MTT, Sigma, USA) was diluted to 0.5mg/ml in cell media and added to the cells. After three hours, MTT solvent (20% (w/v) SDS in 0.02 M HCl) was added to the wells and incubated overnight at 37°C. The next day, absorbance at 570 nm was measured using a plate reader. The background absorbance of MTT without cells was subtracted and the values were normalized to the untreated control condition of each plate, which was set as 100%.

**Amino acid (protein) and nucleotide sequences used in this study:**

**WT α-Synuclein:**
MDVFMKGLSKAKEGVVAAAEKTKQGVAEAAGKTKEGVLYVGSKTKEGVVHGVATVAEK
TKEQVTNVGGAVVTGVTAVAQKTVEGAGSIAAATGFVKKDQLGKNEEGAPQEGILEDMPV
DPDNEAYEMPSEEGYQDYEPEA

**A30P α-Synuclein:**
MDVFMKGLSKAKEGVVAAAEKTKQGVAEA**P**GKTKEGVLYVGSKTKEGVVHGVATVAEK
TKEQVTNVGGAVVTGVTAVAQKTVEGAGSIAAATGFVKKDQLGKNEEGAPQEGILEDMPV
DPDNEAYEMPSEEGYQDYEPEA

**H50Q α-Synuclein:**
MDVFMKGLSKAKEGVVAAAEKTKQGVAEAAGKTKEGVLYVGSKTKEGVV**Q**GVATVAEK
TKEQVTNVGGAVVTGVTAVAQKTVEGAGSIAAATGFVKKDQLGKNEEGAPQEGILEDMPV
DPDNEAYEMPSEEGYQDYEPEA

**G51D α-Synuclein:**
MDVFMKGLSKAKEGVVAAAEKTKQGVAEAAGKTKEGVLYVGSKTKEGVVH**D**VATVAEK
TKEQVTNVGGAVVTGVTAVAQKTVEGAGSIAAATGFVKKDQLGKNEEGAPQEGILEDMPV
DPDNEAYEMPSEEGYQDYEPEA

**A53T α-Synuclein:**
MDVFMKGLSKAKEGVVAAAEKTKQGVAEAAGKTKEGVLYVGSKTKEGVVHGV**T**TVAEK
TKEQVTNVGGAVVTGVTAVAQKTVEGAGSIAAATGFVKKDQLGKNEEGAPQEGILEDMPV
DPDNEAYEMPSEEGYQDYEPEA

**ΔC1-125 α-Synuclein:**
MDVFMKGLSKAKEGVVAAAEKTKQGVAEAAGKTKEGVLYVGSKTKEGVVHGVATVAEK

TKEQVTNVGGAVVTGVTAVAQKTVEGAGSIAAATGFVKKDQLGKNEEGAPQEGILEDMPV
DPDNEAY

**ΔC1-110 α-Synuclein:**
MDVFMKGLSKAKEGVVAAAEKTKQGVAEAAGKTKEGVLYVGSKTKEGVVHGVATVAEK
TKEQVTNVGGAVVTGVTAVAQKTVEGAGSIAAATGFVKKDQLGKNEEGAPQE

**Core α-Synuclein:**
AGKTKEGVLYVGSKTKEGVVHGVATVAEKTKEQVTNVGGAVVTGVTAVAQKTVEGAGSI
AAATGFVKKDQLGKNEEGAPQE

**$C_{rev}$ α-Synuclein:**
MDVFMKGLSKAKEGVVAAAEKTKQGVAEAAGKTKEGVLYVGSKTKEGVVHGVATVAEK
TKEQVTNVGGAVVTGVTAVAQKTVEGAGSIAAATGFVKKDQLGKNEEGAPQEGILE**K**MPV
DP**K**NEAY**K**MPSE**K**GYQDY**K**PEA

**A140C α-Synuclein:**
MDVFMKGLSKAKEGVVAAAEKTKQGVAEAAGKTKEGVLYVGSKTKEGVVHGVATVAEK
TKEQVTNVGGAVVTGVTAVAQKTVEGAGSIAAATGFVKKDQLGKNEEGAPQEGILEDMPV
DPDNEAYEMPSEEGYQDYEPE**C**

**DDX4N1 CtoA:**
GMGDEDWEAEINPHMSSYVPIFEKDRYSGENGDNFNRTPASSSEMDDGPSRRDHFMKSGFA
SGRNFGNRDAGEANKRDNTSTMGGFGVGKSFGNRGFSNSRFEDGDSSGFWRESSNDAEDN
PTRNRGFSKRGGYRDGNNSEASGPYRRGGRGSFRGARGGFGLGSPNNDLDPDEAMQRTGG
LFGSRRPVLSGTGNGDTSQSRSGSGSERGGYKGLNEEVITGSGKNSWKSEAEGGES

**DDX4N1-YFP: His_tag-GST-TEV_site-Ddx4n1-YFP: [HisGSTTEVsiteDdx4n1YFP]**
HHHHHHMSPILGYWKIKGLVQPTRLLLEYLEEKYEEHLYERDEGDKWRNKKFELGLEFPNL
PYYIDGDVKLTQSMAIIRYIADKHNMLGGCPKERAEISMLEGAVLDIRYGVSRIAYSKDFET
LKVDFLSKLPEMLKMFEDRLCHKTYLNGDHVTHPDFMLYDALDVVLYMDPMCLDAFPKL
VCFKKRIEAIPQIDKYLKSSKYIAWPLQGWQATFGGGDHPPKSDLVPRGSPGIHRDENLYFQ
GGAMGSNMGDEDWEAEINPHMSSYVPIFEKDRYSGENGDNFNRTPASSSEMDDGPSRRDH
FMKSGFASGRNFGNRDAGECNKRDNTSTMGGFGVGKSFGNRGFSNSRFEDGDSSGFWRES
SNDCEDNPTRNRGFSKRGGYRDGNNSEASGPYRRGGRGSFRGCRGGFGLGSPNNDLDPDEC
MQRTGGLFGSRRPVLSGTGNGDTSQSRSGSGSERGGYKGLNEEVITGSGKNSWKSEAEGGE
SSDTQGPKVTLQMVSKGEELFTGVVPILVELDGDVNGHKFSVSGEGEGDATYGKLTLKFICT
TGKLPVPWPTLVTTFGYGLMCFARYPDHMKQHDFFKSAMPEGYVQERTIFFKDDGNYKTR
AEVKFEGDTLVNRIELKGIDFKEDGNILGHKLEYNYNSHNVYIMADKQKNGIKVNFKIRHNI
EDGSVQLADHYQQNTPIGDGPVLLPDNHYLSYQSKLSKDPNEKRDHMVLLEFVTAAGIT

**$RP_3$:** RRASL RRASL RRASL

**ssDNA:** 5'-TTT TTC CTA GAG AGT AGA GCC TGC TTC GTG G-3'

**SUPPLEMENTARY FIGURES:**

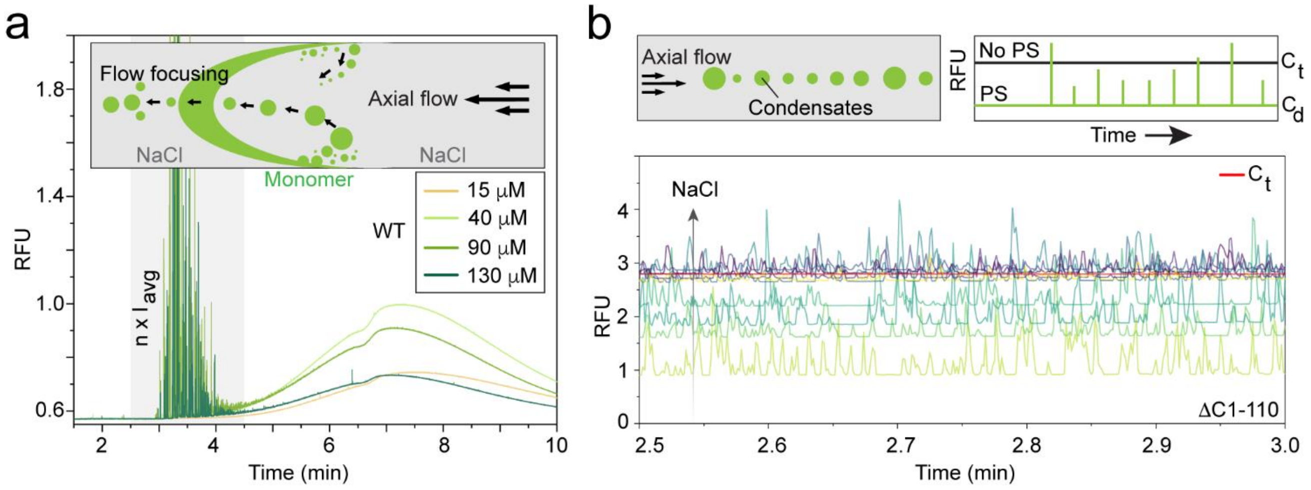


**Supplementary Figure 1: TDIPS and Capflex methods**: **a.** In TDIPS, a small protein plug can be sandwiched between buffer containing an inducing agent for phase separation (for α-Syn, the inducer is NaCl). Diffusion of α-Syn from NaCl depleted regions (plug) to NaCl enriched regions (buffer) induces phase separation at specific locations along the capillary[3]. Larger, more stable condensates are aligned at the center of the capillary and can travel to the front end of the protein plug, appearing as spikes. On the other hand, smaller condensates at the back of the plug can dissolve during their migration—resulting in mass transfer that appears as splitting/shoulder formation of a Taylor-dispersed Gaussian profile of the protein plug. A representative dataset is shown where increasing α-Syn concentrations result in phase separation in TDIPS. The experiment is performed with 25% (w/v) PEG-8000, in 20 mM sodium phosphate buffer (250 mM NaCl), and at 25°C. **b.** In Capflex, a relative decrease in the fluorescence baseline is used to calculate the dilute phase concentration ($C_{dil}$) with respect to the total protein concentration ($C_t$). Condensates appear as individual spikes in a Capflex trace and the heights of the spikes scale with their volume (size)[2]. A representative dataset with 200 μM ΔC1-110 α-Syn showing progressive decrease in $C_{dil}$ along with appearance of spikes, as NaCl concentration decreases. The experiment is performed with 25% (w/v) PEG-8000, in 20 mM sodium phosphate buffer, and at 25°C.

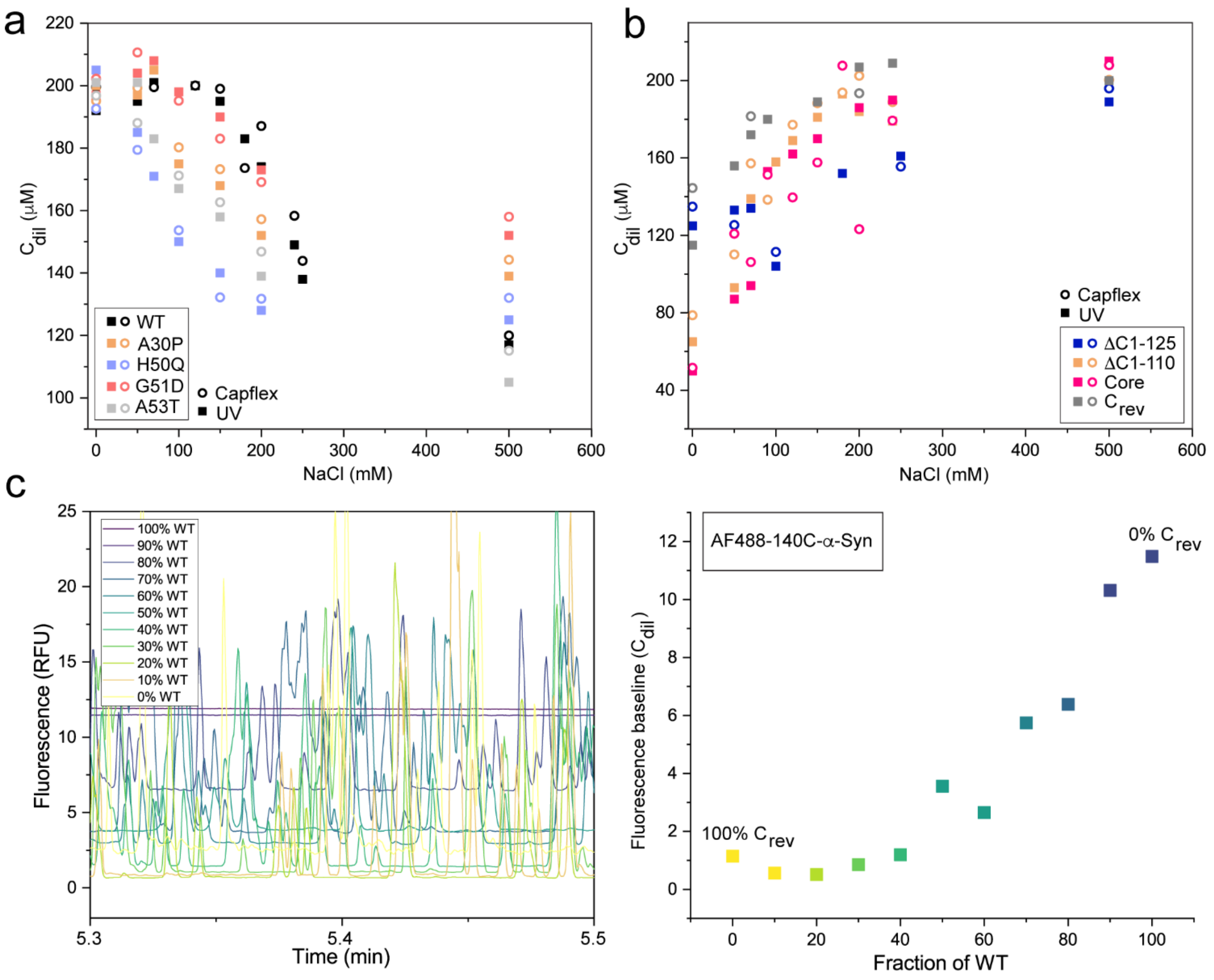


**Supplementary Figure 2: Dilute phase concentrations after phase separation: a.** $C_{dil}$ as a function of NaCl concentration for WT and familial (A30P, H50Q, G51D, A53T) α-Syn variants. **b.** $C_{dil}$ as a function of NaCl concentration for WT and terminally truncated/altered α-Syn variants (ΔC1-125, ΔC1-110, Core, $C_{rev}$). The experiments are performed at $C_t$=200 μM with 20% (w/v) PEG-8000, in 20 mM sodium phosphate buffer, pH 7.4, and at 25°C. The datapoints are collected using UV absorbance (open circles) and Capflex (squares) which are consistent with each other. 50 nM of Alexa488-140C-α-Syn is used as a probe for all Capflex experiments. **c.** (Left) Capflex traces with 25 μM total α-Syn with varying fraction of WT and $C_{rev}$ showing progressive decrease in fluorescence baseline ($C_{dil}$) along with appearance of spikes, as the fraction $C_{rev}$ increases. The experiment is performed with 20% (w/v) PEG-8000, at pH 6.0, in 20 mM sodium phosphate buffer, and at 25°C. (Left) The fluorescence baselines from Capflex is plotted as a function of WT fraction, which indicates partitioning of the Alexa488-140C-α-Syn in a co-phase separating system with both WT and $C_{rev}$ α-Syn.

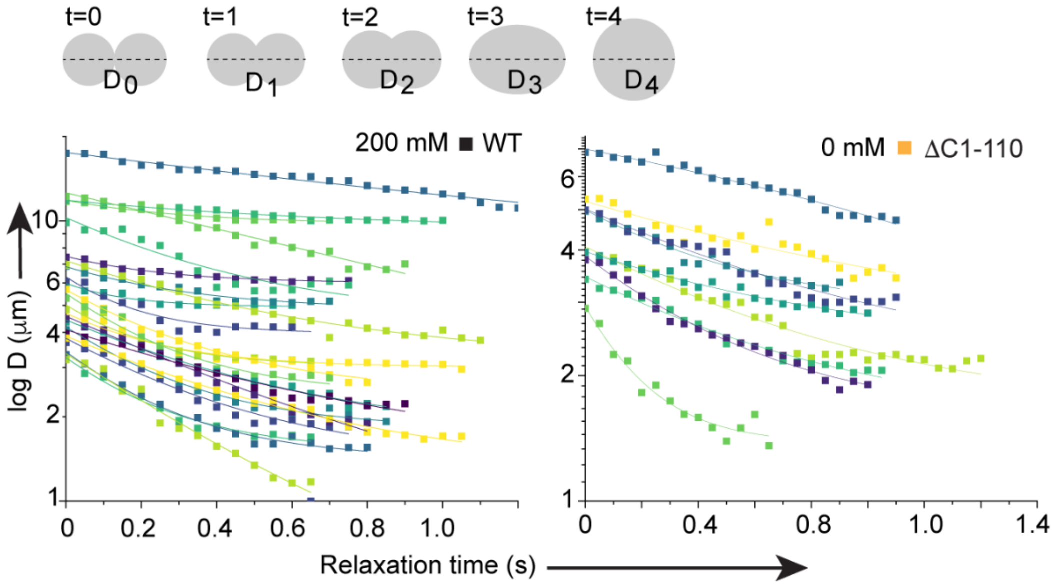


**Supplementary Figure 3: Fusion events detected for WT and ΔC1-110 α-Syn:** Distance (D) as a function of relaxation time (t) (top, schematic) is plotted for fusion events detected for WT (bottom left) and ΔC1-110 (bottom right). The curves are fitted with a mono-exponential decay (see methods) to determine τ values. The experiments are performed with 200 μM α-Syn, 25% (w/v) PEG-8000, in 20 mM sodium phosphate buffer, pH 7.4, and at 25°C. The proteins are phase separated under optimal conditions, i.e., WT at 250 mM NaCl, and ΔC1-110 at 0 mM NaCl. The Y-axis (D) is plotted in log scale for easier visualization.

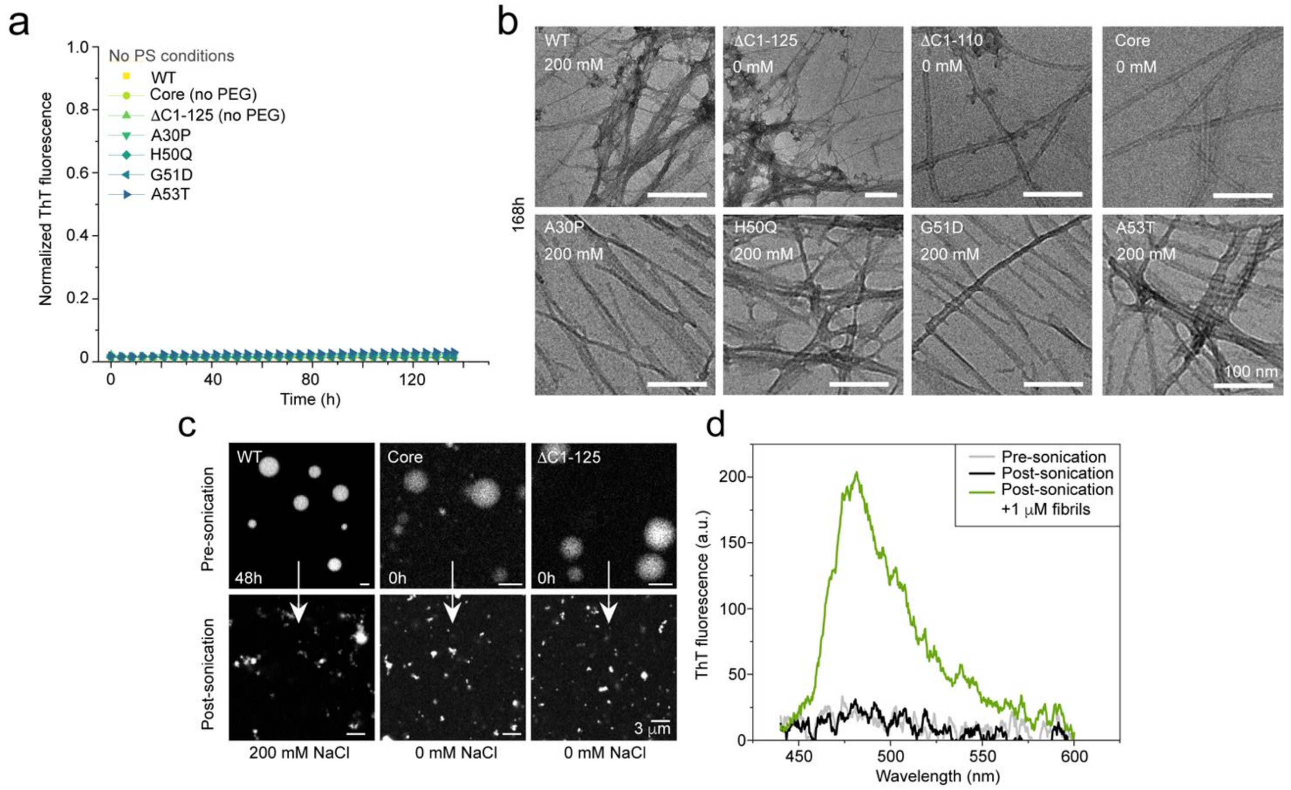


**Supplementary Figure 4: Amyloid aggregation under phase separating conditions: a.** Under soluble (no PS) conditions, none of the α-Syn variants showed aggregation under quiescent conditions, at 25°C, during several days. For full-length variants, the experiment is performed with 200 μM aSyn, 20% (w/v) PEG-8000, in 20 mM sodium phosphate buffer, and without NaCl. Since truncated variants phase separated both in the absence and presence of NaCl, no PEG is added to avoid PS. The rest of the experimental parameters for the truncated variants remain identical to the full-length variants. **b.** Representative TEM images of α-Syn variants under optimal phase separating conditions after 7 days showing the presence of amyloid fibrils. The samples are phase separated under optimal conditions, and aged (at 25°C) without shaking. The samples were prepared with 200 μM α-Syn, 20% (w/v) PEG-8000, in 20 mM sodium phosphate buffer, pH 7.4. **c.** Representative confocal fluorescence images showing sonication can break apart aggregated condensates of WT (sonication time: 48h), ΔC1-110 and Core (sonication time: 0h). The condensates are probed with 100 nM Alexa488-140C-α-Syn. **d.** A 200 μM monomeric α-Syn solution (no PEG) in 20 mM sodium phosphate buffer with 200 μM ThT (pH 7.4), is sonicated to check whether sonication alone could induce aggregation. To ensure whether ThT is active post-sonication, 1 μM amyloid fibrils are added to the same sample.

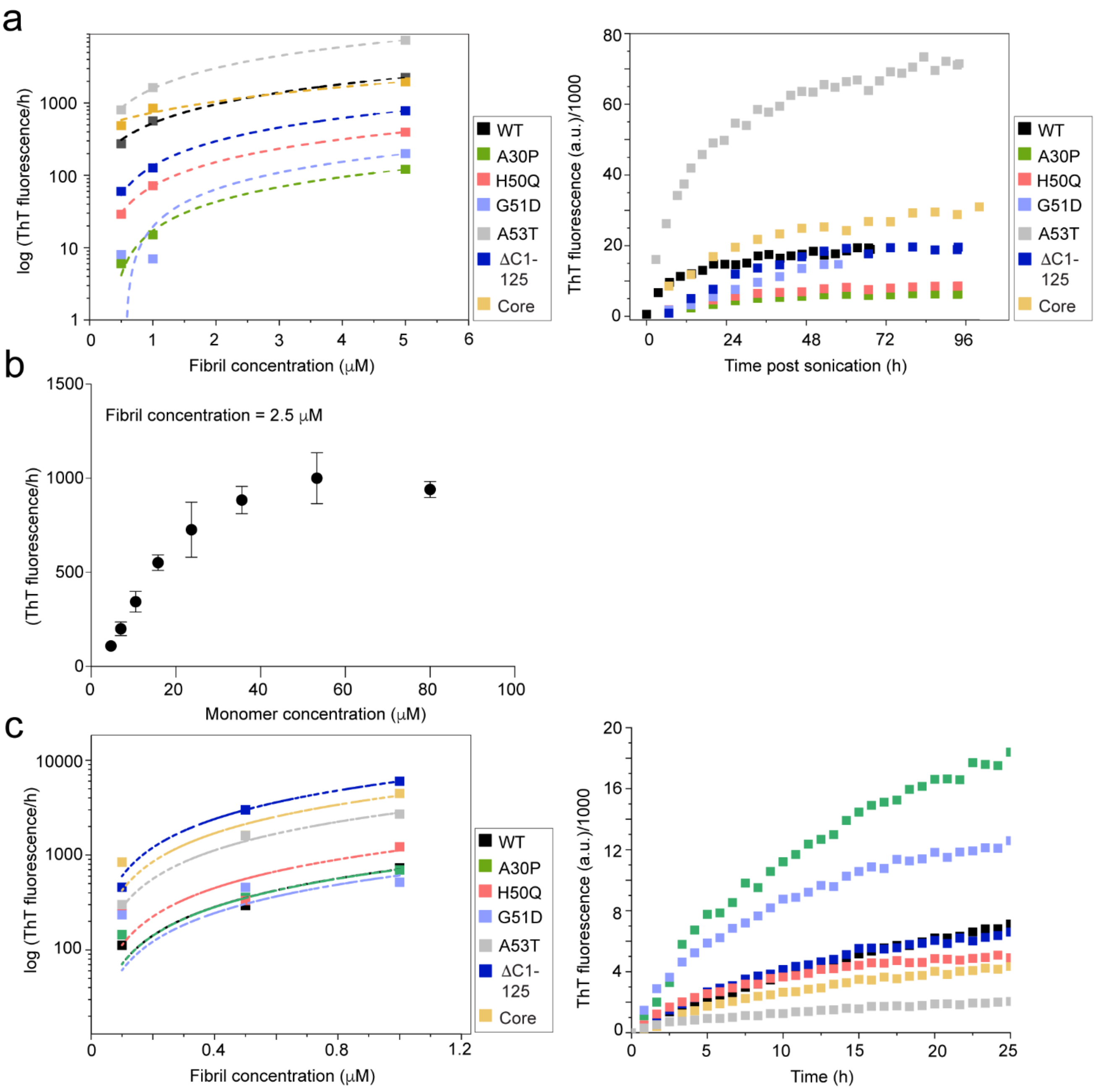


**Supplementary Figure 5: Estimation of templating competent amyloid fibrils formed within α-Syn condensates: a.** (Left) Calibration experiment to calculate the elongation rate constant ($k_+$, slope of the fitted line) of preformed amyloid fibrils for different α-Syn variants. The datapoints are fitted with a linear equation and plotted in a log-scale for easier visualization. The calibration experiments are performed under quiescent conditions, with 30 μM monomeric α-Syn, with 15% (w/v) PEG-8000, in 20 mM sodium phosphate buffer with 30 μM ThT (pH 7.4). (Right) Elongation profiles of representative set of WT and variant α-Syn condensate solutions post sonication (48h for full-length and 0h for truncated variants). The initial ThT fluorescence value is normalized to zero. The experiments are performed under quiescent conditions, with 20% (w/v) PEG-8000, in 20 mM sodium phosphate buffer, pH 7.4, and at 25°C. Optimal phase separating conditions are chosen for each variant. Initial datapoints (4-5h post sonication) from these ThT traces are used to calculate the respective elongation rates. **b.** Elongation rates (ThT/h) as a function of WT monomer concentration. The experiment is performed in presence of 2.5 μM WT α-Syn fibrils, under quiescent conditions, and at 25°C. Values represent mean±S.D for n=3 independent experiments. **c.** (Left) Calibration experiment to calculate the elongation rate constant ($k_+$, slope of the fitted line) of preformed amyloid fibrils for different α-Syn variants. The datapoints are fitted with a linear equation and plotted in a log-scale for easier visualization. The calibration experiments are performed under quiescent conditions, with 30 μM monomeric α-Syn, in 20 mM sodium phosphate buffer with 30 μM ThT (pH 7.4). (Right) Elongation profiles of representative set of WT and variant α-Syn condensate solutions post centrifugation, and sonication (isolated at 48h for full-length and 0h for truncated variants). The initial ThT fluorescence value is normalized to zero. The experiments are performed under quiescent conditions, in 20 mM sodium phosphate buffer, pH 7.4, and at 25°C. Initial datapoints (4-5h post sonication) from these ThT traces are used to calculate the respective elongation rates.

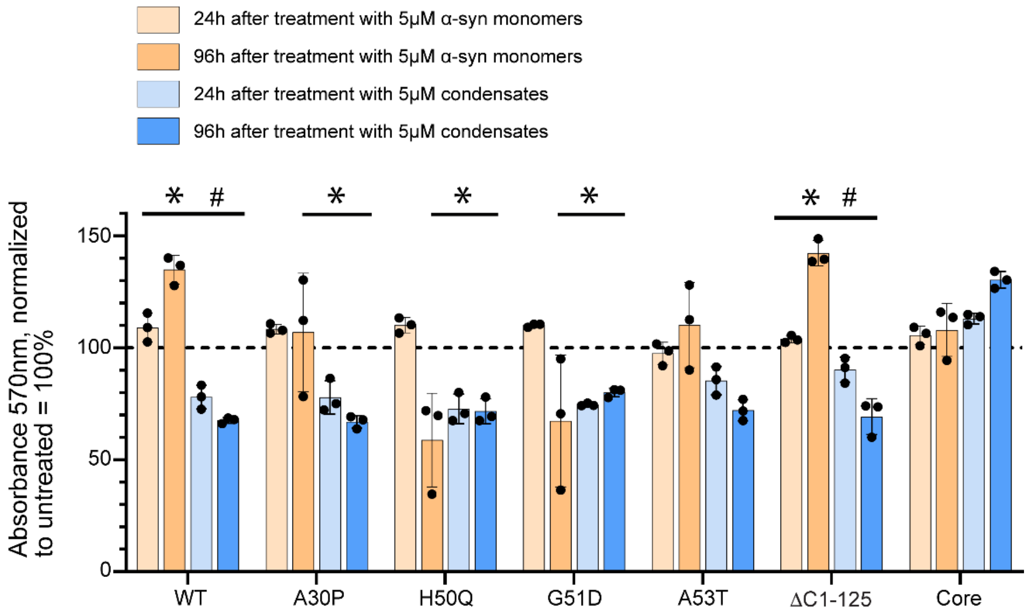


**Supplementary Figure 6: MTT viability assay of SHSY-5Y cells:** Undifferentiated SHSY-5Y cells are treated with 5 µM of monomeric α-Syn and 5 µM monomer-equivalent condensate solution (aged for 7 days) (both WT and variants). After 24 and 96h, cell viability is measured using MTT assay (absorbance at 570 nm). The data is compared to the untreated condition (normalized to 100% viability). Values represent mean±S.D for n=3 independent experiments. Statistical significance is calculated with an unpaired t test, comparing monomers and condensates of each variant at the two indicated time-points. The p-values≤0.05 at 24h are indicated with * (asterisk) and the p-values≤0.05 at 96h are indicated with # (hashtag).

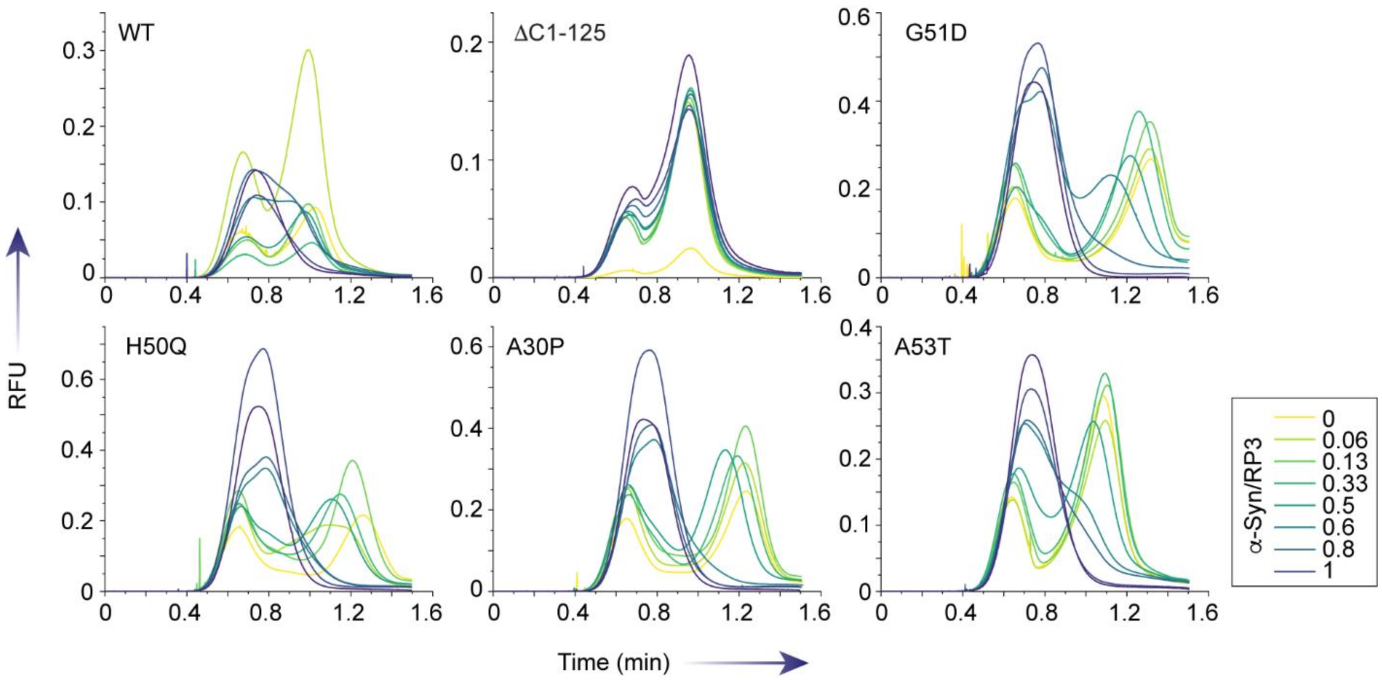


**Supplementary Figure 7: Triple-plug TDIPS for RP3/ssDNA coacervation in the presence of α-Syn variants**: TDIPS showing the effects of α-Syn variants on RP3/ssDNA coacervation. In the absence of α-Syn, RP3 and ssDNA form coacervates, which is detected as splitting (mass transfer) of an otherwise Gaussian profile. Increasing α-Syn concentrations in the ssDNA plugs dissolve coacervates (from two peaks to one peak) in a dose-dependent manner, which is not observed for ΔC1-125 due to the lack of C-terminal acidic (negatively charged) residues. The experiments are performed in 10 mM Tris-HCl, pH 8.0, 150 mM NaCl, and at 20°C. RP3 and ssDNA concentrations in the plugs are 1.5 mM and 10 μM, respectively. α-Syn is added to the ssDNA plug at concentrations ranging from 0-1.5 mM. Molar ratios of α-Syn/RP3 are shown in the legend.

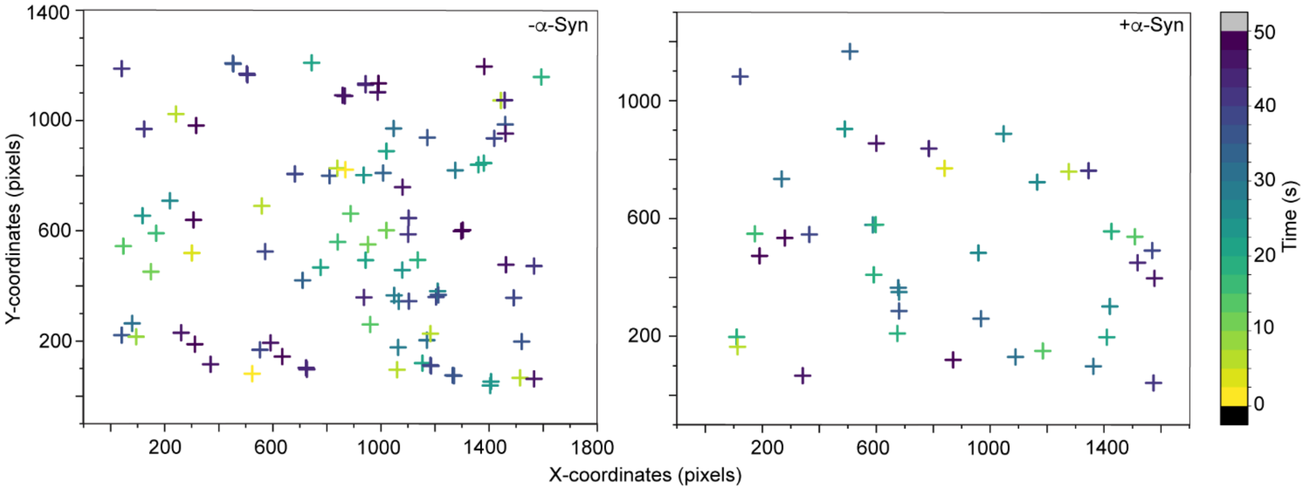


**Supplementary Figure 8: α-Syn restricts DDX4 condensate fusion**: Detected fusion events over 1 min are marked on a microscopic field of view (FOV) spanning 1400x1400 pixels. The samples analyzed are DDX4 condensates alone (left) and DDX4 condensates with 5 μM α-Syn (right). Each coordinate represents one successful fusion event. The color code indicates time of fusion. The experiment is performed n=2 independent times with similar observations. DDX4 is phase separated at 70 μM protein concentration, in 10 mM sodium phosphate buffer, pH 6.5, 50 mM NaCl, and at 25°C.

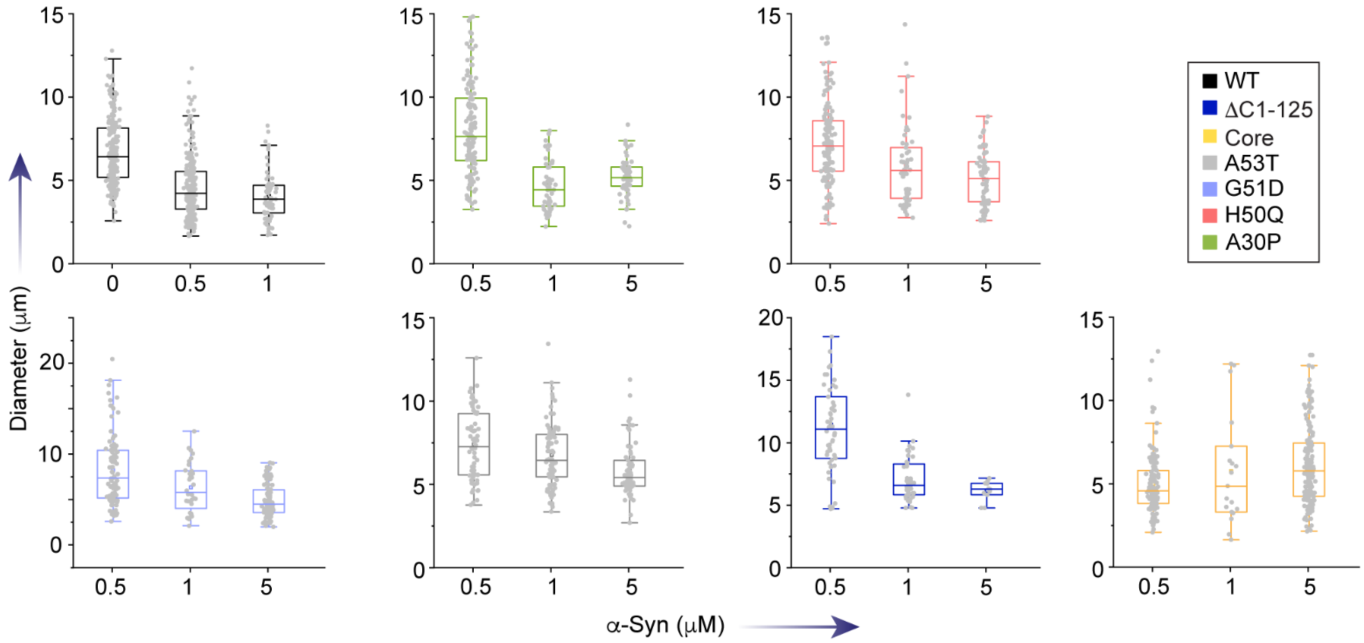


**Supplementary Figure 9: Size distribution of DDX4 condensates in presence of α-Syn**: Distribution of DDX4 condensate diameters in the presence of increasing α-Syn (WT and variant) concentrations (sub-stoichiometric, 0-5 μM) showing a narrowing of the distribution with α-Syn with a progressive decrease in the average condensate diameter. This effect is least prominent for Core α-Syn. DDX4 is phase separated at 70 μM protein concentration, in 10 mM sodium phosphate buffer, pH 6.5, 50 mM NaCl, and at 25°C.

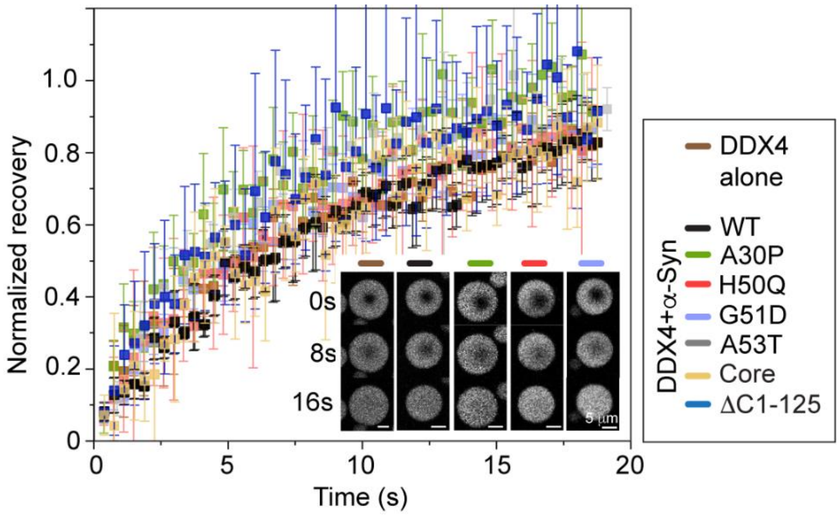


**Supplementary Figure 10: FRAP on DDX4 condensates in presence of α-Syn:** (Left) Normalized recovery of DDX4-YFP molecules within DDX4 condensates in presence of 5 μM WT and variant α-Syn obtained from FRAP experiments. Values represent mean±SD for n=3 independent experiments (obtained from a total of 5-6 condensates for each variant). (Inset) Confocal microscopy images of DDX4 condensates alone, and in presence of some of the representative variants: 5 μM WT, A30P, H50Q, and G51D α-Syn at 0s, 10s and 20s post bleach are shown. The experiments are performed with 70 μM DDX4 (100 nM DDX4-YFP as fluorescent reporter), in 10 mM sodium phosphate buffer, pH 6.5, 50 mM NaCl, and at 25°C.

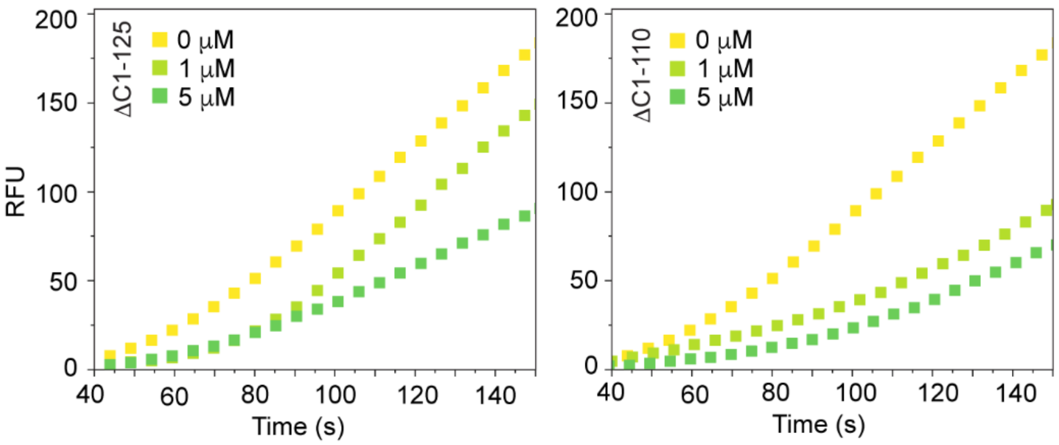


**Supplementary Figure 11: Partitioning of ssDNA within DDX4 condensates in presence of truncated α-Syn:** Background corrected fluorescence intensity of Alexa488-ssDNA within DDX4 condensates is plotted in absence and presence of 1 and 5 μM truncated (ΔC1-125 and ΔC1-110) α-Syn variants. The slopes (rates, RFU/s) are calculated by fitting the datapoints with a linear equation. The datapoints represent mean values obtained from N=50 condensates. The experiments are performed with 70 μM DDX4, in 10 mM sodium phosphate buffer, pH 6.5, 50 mM NaCl, and at 25°C.
